# Hemifusomes and Interacting Proteolipid Nanodroplets: Formation of a Novel Cellular Organelle Complex

**DOI:** 10.1101/2024.08.28.610112

**Authors:** Amirrasoul Tavakoli, Shiqiong Hu, Seham Ebrahim, Bechara Kachar

## Abstract

Within cells, vesicle fusion, scission, and the formation of intraluminal vesicles are critical processes that facilitate traffic, degradation, and recycling of cellular components, and maintenance of cellular homeostasis. Despite significant advancements in elucidating the molecular mechanisms that drive these dynamic processes, the direct in situ visualization of membrane remodeling intermediates remains challenging. Here, through the application of cryo-electron tomography in mammalian cells, we have identified a previously undescribed vesicular organelle complex with unique membrane topology: heterotypic hemifused vesicles that share extended hemifusion diaphragms (HDs) with a 42 nm proteolipid nanodroplet (PND) at their rim. We have termed these organelle complexes “hemifusomes”. The HDs of hemifusomes exhibit a range of sizes and curvatures, including the formation of lens-shaped compartments encapsulated within the membrane bilayer. The morphological diversity of the lens-shaped vesicle aligns with a step-wise process of their intraluminal budding, ultimately leading to their scission and the generation of intraluminal vesicles. We propose that hemifusomes function as versatile platforms for protein and lipid sorting and as central hubs for the biogenesis of intraluminal vesicles and the formation of multivesicular bodies.

## Introduction

The endolysosomal and secretory systems-comprising networks of structurally and functionally diverse membrane-bound organelles undergoing constant remodeling, trafficking, and recycling or degradation-are integral to cellular function and homeostasis. Central to this broad range of cellular activities are membrane fusion events required for membrane and content mixing, and membrane budding, and scission involved in the genesis of intraluminal vesicles ^1–4^. Despite significant research elucidating the molecular machineries involved in these processes ^5–7^, the direct, in-situ visualization and characterization of intermediate structures of membrane fusion and scission remain formidable challenges ^8^. Similarly, while extensive literature exists on the protein machineries required for the formation of intraluminal vesicles, the structural intermediates of membrane budding and scission events that occur during this process remain poorly documented and inadequately understood ^5,7,9^. The precise mechanisms by which membrane remodeling is coordinated to ensure the proper trafficking and turnover of membrane proteins and lipids and to maintain cellular health and function remain topics of broad interest and vigorous research. Dysregulation of secretory and endolysosomal pathways has been implicated in numerous diseases ^10–12^, underscoring the importance of understanding these processes for the development of targeted therapies.

Recent advances in cryo-electron tomography (cryo-ET) have enabled unprecedented visualization of cellular structures in near-native states ^13–17^, yielding novel insights into cellular structure/function relationships. The current study leverages in situ cryo-ET to uncover a novel class of organelles in mammalian cells, termed ‘hemifusomes’. These organelles, found in all four different cell types we investigated, are characterized by heterotypic vesicle pairs hemifused via expanded hemifusion diaphragms (HDs), a unique membrane topology previously presumed to be too unstable for any biological function beyond serving as fleeting intermediates in membrane fusion and scission ^18–20^.

Hemifusomes appear in two related morphological configurations: (1) a direct hemifusion of two heterotypic vesicles; and (2) a flipped conformation where an intraluminal vesicle is hemifused to the luminal side of the membrane bilayer of a larger vesicle. In both cases, a smaller vesicle and a larger vesicle share a HD. In the case of direct hemifusomes, the topology and content of the paired heterotypic vesicles are strikingly consistent. Additionally consistent is the association of a proteolipid particle or droplet with the rim of the HD. In the case of flipped hemifusomes, more complex compound fusion configurations often arise. We explore relationships between these morphologically diverse but topologically similar conformations and their potential participation in endolysosomal functions and dynamics, including formation of intraluminal vesicles and biogenesis of multivesicular bodies.

## Results

### Observation of hemifused vesicles at the periphery of cultured cells

The periphery of vitrified mammalian cells (COS-7, HeLa, Rat-1, and 3T3 cells) of approximately 250 to 450 nm ice thickness, were surveyed using a Krios (Thermo-Fisher) cryo-electron microscope operating at 300 kV (Fig. S1a and b). These thin cellular regions enabled visualization of membrane-bound organelles that were restricted to diameters of up to 400 nm (Figs. 1a, S1, and S2). During a lower magnification initial survey of this region, we identified vesicles that, based on size, morphology and content, are likely to be endosomes, lysosomes, or MVBs (Figs. 1 and S1). While endosomes and lysosomes at the leading edge of the cell were predominantly spherical (Fig. 1 and 2), we did observe features such as tubulation and budding, characteristic of larger endosomes and lysosomes, further away from the cell periphery (Fig. S2 b and e).

**Figure 1.**
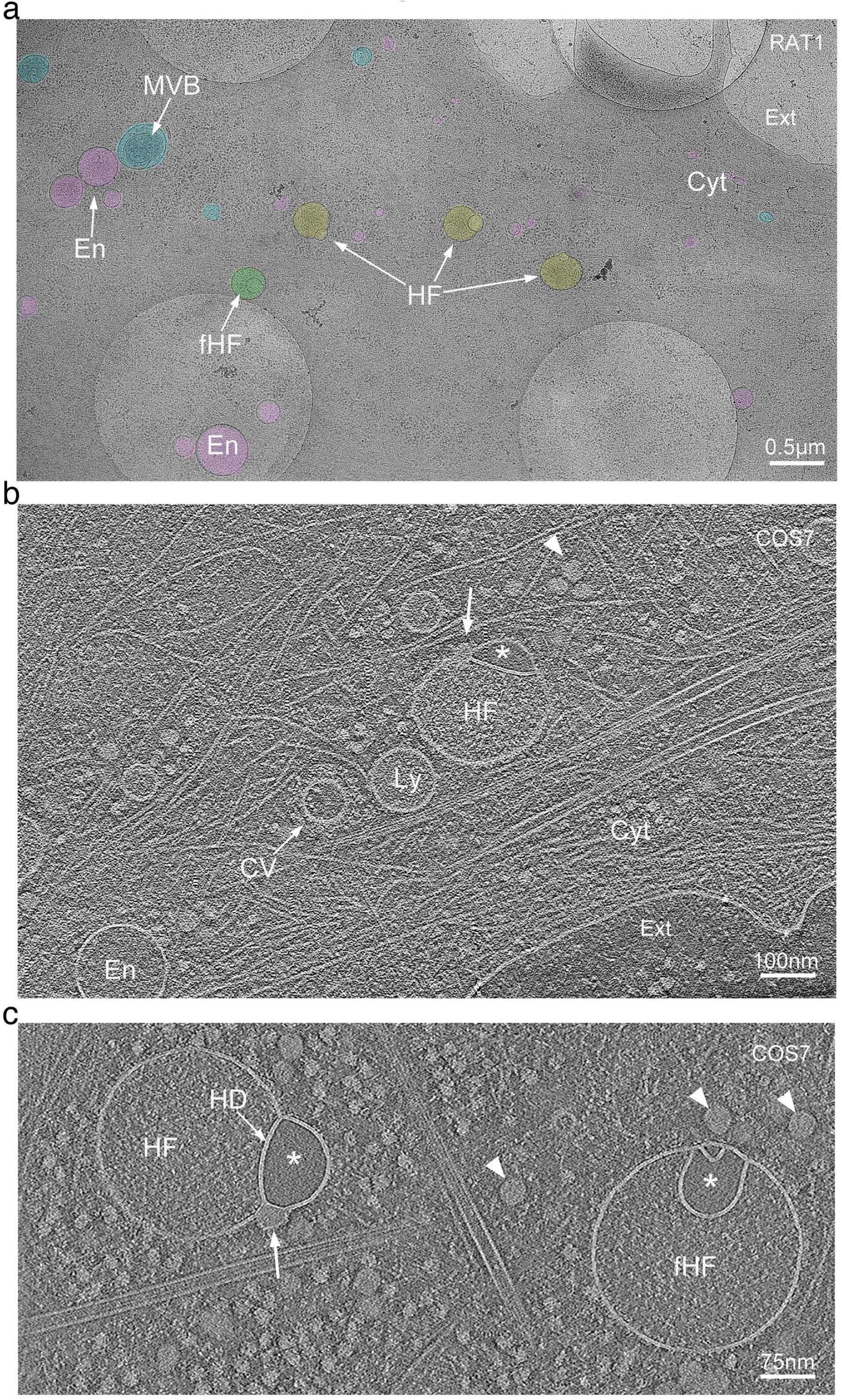
Cryo-electron tomography observation of hemifused vesicles at the leading edge of cultured cells. a. Representative cryo-electron microscopy image of the leading edge of a RAT1 cell cultured on cryo-EM grids. Lamellipodia and filopodia in the upper right corner delineate the cell border, separating the cytoplasm (Cyt) from the extracellular space (Ext). Vesicular organelles are highlighted in color: endosomes (En, pink), multivesicular bodies (MVB, blue), hemifusomes (HF, yellow), and flipped hemifusomes (fHF, green). Scale bar: 0.5 µm. b. Representative cryo-electron tomogram slice of the border of a COS7 cell highlighting cytoskeletal components, endosomes (En), lysosomes (Ly), a clathrin-coated vesicle (CV), and a hemifusome (HF). A single bilayer or hemifusion diaphragm separates the hemifused vesicles. The larger vesicle has a fine granular content, and the smaller hemifused vesicle has a smooth translucent lumen (*). The arrowhead points to the proteolipid particle and arrow points to the nanodroplet at the rim of the hemifusion diaphragm. Scale bar: 100 nm. c. Tomographic mid-cross-section through a direct (HF) and a flipped hemifusome (fHF) showing the well-defined bilayer outline of the vesicle membranes. A single bilayer or hemifusion diaphragm (HD) separates the hemifused vesicles. Arrowheads point to the proteolipid particle, as well as similar proteo-lipid particles seen free in the cytoplasm. Scale bar: 75 nm.

**Figure 2.**
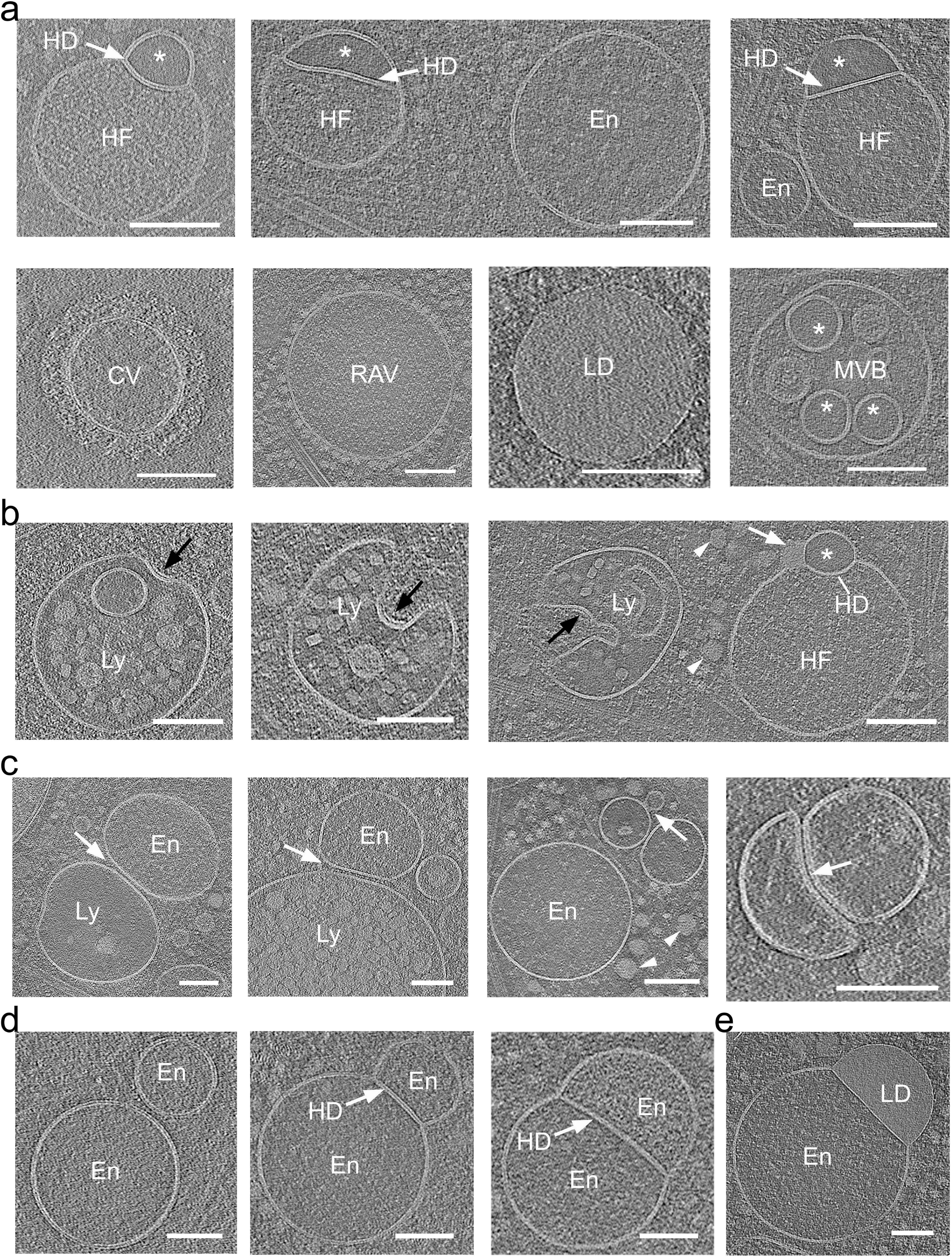
Hemifusome luminal content is distinct from other membrane-bound organelles of the endo-lysosomal system. a. Representative cryo-electron microscopy images of various membrane-bound organelles within the endo-lysosomal system. Endosomes (En), lysosomes (Ly), a clathrin-coated vesicle (CV), ribosome associated vesicles (RAV), multivesicular body (MVB), lipid droplet (LD), and hemifusomes (HF) are identified. The distinct luminal content of each organelle is visible, with the smaller hemifusome vesicles (*) consistently showing a unique light, smooth, and particle-free luminal content compared to other organelles. Similar smooth lumen vesicles were found only inside some MVBs (*). Hemifusion diaphragm (HD) is highlighted with white arrows. b. Series of lysosomes at various initial stages of inward budding of a vesicle obtained from tomographic slices. The last panel shows a side-by-side comparison of a lysosome and a hemifusome (HF). Black arrows point to a distinct surface protein complex, likely ESCRT, at the inwardly curved portion of the lysosomal membrane. White arrowhead points to proteolipid nanodroplets in the cytoplasm and white arrow points to the nanodroplet at the rim of the hemifusion diaphragm. c. Series of tomogram slices showing endosomal (En) and lysosomal (Ly) vesicles adhered or docked to each other (white arrows). d. Series of tomogram slices showing hemifused endosomes sharing an extended hemifusion diaphragm (HD) (white arrow). Right panel-a hemifused endosome (En) and lipid droplet (LD) Note the granular texture of the lumen of all vesicles. Scale bars: 100 nm.

Strikingly, we also identified closely interacting pairs of vesicles in two previously undescribed conformations: the first, a smaller vesicle appearing hemifused to the cytoplasmic side of a larger vesicle, and the second, a flipped configuration, where an intraluminal vesicle appears hemifused with the luminal side of the larger vesicle (Figs. 1 and S1). These hemifused organelles were observed in all four cell types examined (Figs. 1a and S1b, d, e). Surveying randomly selected regions from the periphery of COS-7 cells (∼10 µm^2^ search projections as in Fig. S1b, n=81), we estimated the frequency of these hemifused vesicles to be approximately 10% of the total number of membrane-bound organelles (average number of hemifused vesicles= 0.6 +/− 0.7; average number of other vesicles= 6.6 +/− 4.0), with broad local variation within and between cells (Figs. 1 and S1).

### Cryo-electron tomography of hemifused vesicles

For a more detailed analysis of morphology of the hemifused vesicles and their spatial context within the cytoplasm, we performed low-dose tilt series image acquisitions for cryo-ET, following the methodology described by Hagen et al. (2017). The workflow for this procedure is detailed in the Methods section, and representative images are shown in Fig. S1a-c. Tomogram slices of the periphery of the cell revealed cytoskeletal components and more detailed views of vesicular elements, including clathrin-coated vesicles, endosomes, lysosomes, MVBs and the previously undescribed hemifused vesicles (Figs. 1b and c). Tomographic mid-cross-sections through direct and flipped hemifusomes revealed the well-defined bilayer outline of the vesicle membranes (Figs. 1b, c and S1c). The tomograms also confirmed the presence of a shared bilayer at the interface of the vesicle pair (Figs. 1b, c and S1c), characteristic of an extended HD, a fusion intermediate theorized ^18–21^ and observed in synthetic ^22^ or cell-free ^8,14,23^ systems, but rarely observed in intact cells ^24,25^.

Based on their size, electron density, and subcellular localization (Fig.1 and 2 and S1 and S2), often found near endosomes (Fig. S2), we speculate that the hemifused vesicle complexes are either a new class of organelle, or previously unrecognized intermediates or members of the broader endo-lysosomal system. Of note, while the lumen of the larger vesicles displayed a fine granular texture, comparable to the lumen of endosomes or early lysosomes, the smaller vesicles in these pairs consistently exhibited “translucent” content devoid of visible differential electron scattering or phase contrast typical for granular or particulate material (Figs. 1b and c, and Figs. 2 and 3), we posit that the translucent luminal content likely reflects a protein-free or very dilute aqueous solutions.

**Figure 3.**
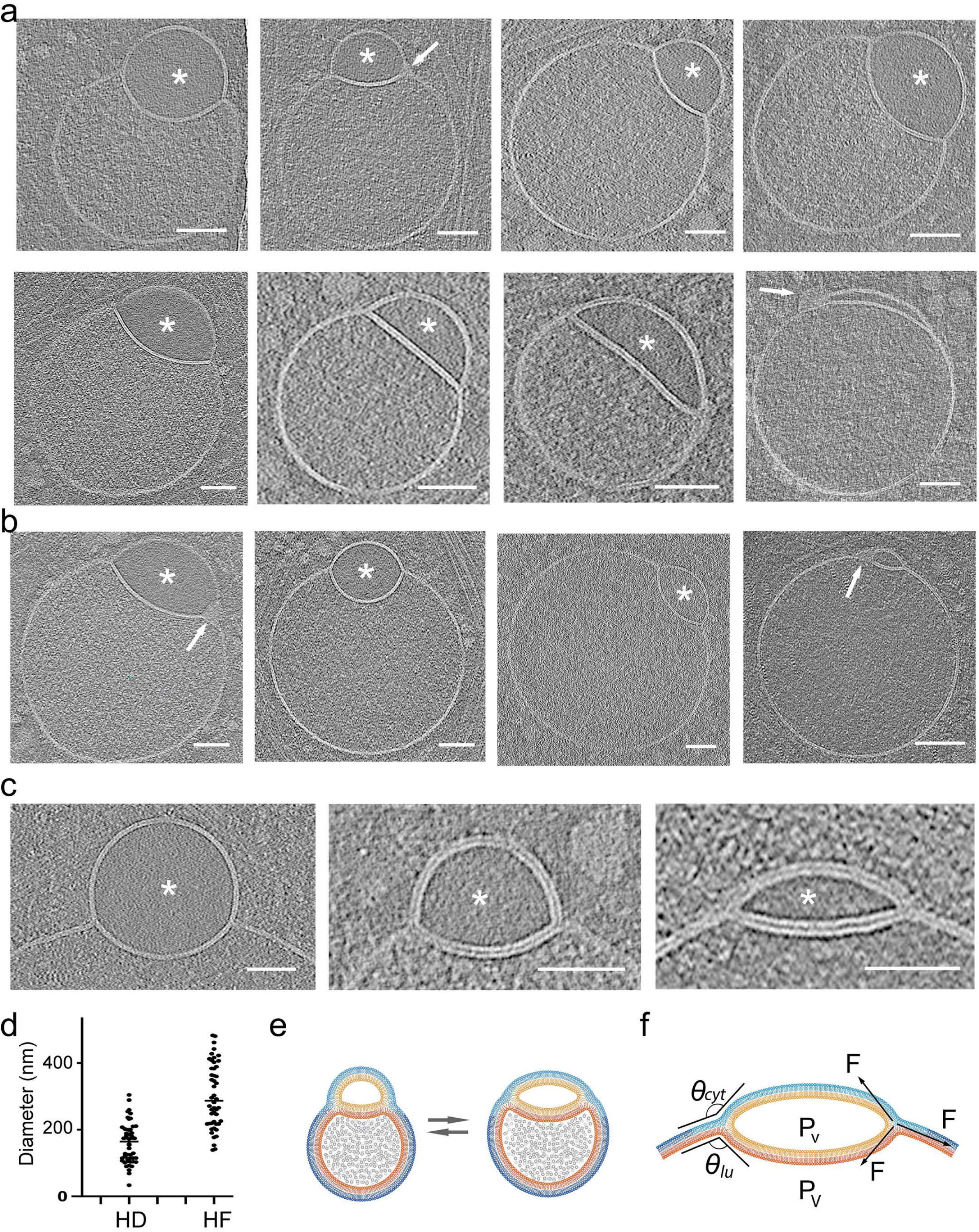
Range of morphological appearances of direct hemifusomes. a. Cryo-electron tomography mid-cross-section slices of various direct hemifusomes highlighting the variability in sizes and shapes across and within each hemifusome pair. Note the smooth appearance of the smaller vesicle of the hemifusome (*). Hemifusomes also show variability in their hemifusion diaphragm diameter and curvature. Arrows point to proteolipid particles lodged in the hemifusome at the rim of the hemifusion diaphragm where it meets with membranes of the two vesicles. b. Hemifusomes showing deformation of the smaller vesicle with the expansion and curvature of the hemifusion diaphragm, resulting in a cross-sectional view that resembles a lens-shaped vesicle. In this specific configuration, the entire inner leaflet as well as the content of the smaller vesicle becomes fully embedded within the bilayer of the membrane containing the larger vesicle. c. Close-up views of the hemifusion diaphragm and degrees of flattening of the smaller vesicle (*) to form the lens-shaped structure embedded in the bilayer. d. Diameters of the hemifusion diaphragm (HD), mean= 158.4 +/− 60.9, and the larger vesicle of the hemifusome (HF), mean= 299.3 +/− 96.2. n = 50 hemifusomes. e. Diagram illustrating the hemifused configuration and possible interconversion between hemifusome conformations. f. Diagram illustrating some of the forces at play to modulate hemifusome angles between the membranes and overall shape. θ = angle of the cytoplasmic (*cyt*) and luminal (*lu*) leaflets of the bilayer at the point of junction with the hemifusion diaphragm. ; P, internal pressure of smaller (v) and larger vesicle (V), F, membrane tension vectors.

We designate the term “direct hemifusomes” or simply “hemifusomes” to the hemifused vesicles where the smaller translucent vesicle is on the cytoplasmic side of the membrane of the larger vesicle, and “flipped hemifusomes” to the conformation where the translucent vesicle is hemifused to the luminal or exoplasmic side of the larger vesicle membrane (Fig.1c). Additionally, 3D tomogram reconstructions frequently revealed a dense or phase-dark particle, or nanodroplet, integrated into the bilayer at the margin of the HD (arrows in Fig. 1b and c). Seemingly membrane-less particles of similar size and overall appearance can be observed in the cytoplasm surrounding the hemifusomes in the tomogram slices (indicated by arrowheads in Fig. 1b, c).

Our observation of hemifusomes in four different cell lines originating from various species and tissues and frozen as close as possible to their native state suggests that they may be common components of the cell periphery in a wide range of cells and tissues. Additionally, a review of archival transmission electron microscopy images of plastic-embedded thin sections of conventionally prepared inner ear tissue from our lab revealed structures resembling hemifused vesicles within the endosomal compartment of epithelial cells (Fig. S2b).

### Comparison of the hemifusome luminal content to the content of other membrane-bound organelles

To explore the nature of the direct hemifusome, we compared its morphology and the appearance of its luminal content with other known vesicular organelles in the surrounding cytoplasm (Fig. 2 a and b). As stated above, the lumen of the larger vesicles in the heterotypic hemifused pair displayed a fine granular texture, comparable to the lumen of endosomes (Fig. 2 a and b). However, the unique smooth and translucent appearance of the luminal content of the smaller vesicle (asterisks in Fig. 2a and b) did not match the texture or electron density of the lumen of any of the other vesicular organelles, including endosomes, clathrin-coated vesicles, ribosome-associated vesicles, lipid droplets (Fig. 2a), or various conformations of lysosomes (Fig. 2b). The only vesicles in the 308 tomograms we acquired with comparable content were some of the intraluminal vesicles in MVBs (asterisks in Fig. 2a, lower panel).

We also observed (at least 10 examples in 308 tomograms) endosomes, lysosomes and lipid droplets, either docked (Fig. 2c) or hemifused (Fig. 2d), in line with established models of docking ^22,26–28^ and fusion that lead to the formation of late endosomes or the delivery of endosomal cargo to lysosomes ^4,29^. Notably, we did not observe any translucent vesicles that were either free or docked to endosomes. Further, we also identified hemifusomes in cellular regions that were largely devoid of membrane-bound organelles (Fig. S1f), where the chances of vesicle encounter, docking, and subsequent fusion to form a hemifused pair would be exceedingly low. Together, these findings led us to question whether hemifusomes might be formed by alternative mechanisms to canonical vesicle fusion.

### Morphological variation of direct hemifusomes

Representative close-up views of the rim of the hemifusome HD, where the bilayers of the two vesicles and the HD meet, enabled us to confirm that the cytoplasmic leaflets of the bilayers of the two interacting vesicles were contiguous (Figs. 1c, 2 and 3). The leaflets of the HD bilayer is comprised of the exoplasmic leaflets of the two interacting vesicles, as expected for hemifused vesicles. These bilayer arrangements are depicted in Fig. 3d. Measurements of the thickness of the bilayer of each hemifusome vesicle, and the shared HD, were comparable to typical membranes at ∼ 4 nm (Fig. S4 g and h). The paired vesicles forming the hemifusome can vary widely in both individual and relative sizes. Additionally, even when the paired vesicles are of comparable-size, their HDs exhibit variability in radius and curvature indicating diverse degrees of radial expansion of the area of hemifusion between vesicles (Fig. 3 and S4).

Independent of how the hemifusomes are formed, radial expansion of HDs must require adhesive forces capable of deforming vesicles and overcoming the ensuing increase of internal pressure and membrane tension as the surface-to-volume ratio changes for both vesicles. The observation of the HD bulging into the larger vesicle is consistent with intrinsic pressure differential that occurs when the HD expands; namely, the smaller vesicle experiences higher pressure due to the faster rate of change in surface-to-volume ratio. However, intriguingly, HDs could also be found, albeit less frequently, with no curvature or even bulging towards the smaller vesicle (Fig. 3a). Local changes in the composition and biophysical properties of the bilayers are expected as the angle/curvature of the monolayers change at the rim of the HD ^21,30–32^.

In addition to the biophysical properties of the membrane ^20^ and other factors intrinsic to the hemifusome complex, the shape of the complex could also be influenced by local physical constraints. We observed hemifusomes deformed by surrounding actin filaments and microtubules (Fig. S3a and b) or by constraints imposed by proximity to the plasma membrane (Fig. S3b-c) as the hemifusome squeezed through the thinner regions at the cell’s edge. These varying conformations illustrate the compressibility and deformability of the hemifusomes and how internal and external forces or constraints (Figs. 3f and S3) potentially contribute to their overall shape, expansion of the HDs, and the angles and curvature of the interacting membranes at the rim of the HDs.

### Emergence of intramembrane lens-shaped structures in hemifusomes

The average radius for the larger vesicle in the hemifusome in the areas we surveyed was measured to be ∼ 299.3 +/− 96.2 nm (n = 50), and for the HDs of the same hemifusomes was measured to be 158.4 +/− 60.9 nm (n = 50) (Fig. 3d). This is an order of magnitude larger than the transient ∼10 nm HD estimated to exist in canonical membrane fusion events ^14,18,33,34^. This finding implies that the expansion of the hemifusome HD is energetically favorable, and large HDs (ratio of HD to hemifusome diameters is shown in Fig. S4) are likely stable with an extended lifespan. At this size, the HDs are still smaller than the diffraction limit of light, explaining why hemifusomes have thus far not been detected by light-microscopy.

Indeed, we often observe hemifusome HDs comparable in size to the remaining membrane of the smaller vesicle (Fig. 3b). In this configuration, the entire inner leaflet and contents of the smaller vesicle are fully encapsulated within the bilayer of the membrane encompassing the larger vesicle. This symmetric intra-bilayer structure has been identified in silico simulations as a long-lived, lens-shaped product of hemifusion between a lipid vesicle and a planar lipid bilayer, referred to as ‘dead-end’ hemifusion ^18^. It is particularly exciting that we observe such long-lived, lens-shaped structures within a complex biological membranous organelle. Consistent with the in-silico simulations, this lens-shaped conformation is observed in hemifusomes containing very small, translucent vesicles (Fig. 3b-c), where the larger vesicle closely approximates the flatness of a planar lipid bilayer.

### Parallels between direct and flipped hemifusomes

In our 308 tomograms, we were able to identify clearly 88 direct hemifusomes (representative examples in Fig. 3) and 48 hemifusomes in the flipped configuration (representative examples in Fig. 4). Based on the conserved topological features of direct and flipped hemifusomes, we hypothesize that they may represent different conformations of the same organelle, potentially transitioning from one form to the other.

**Figure 4.**
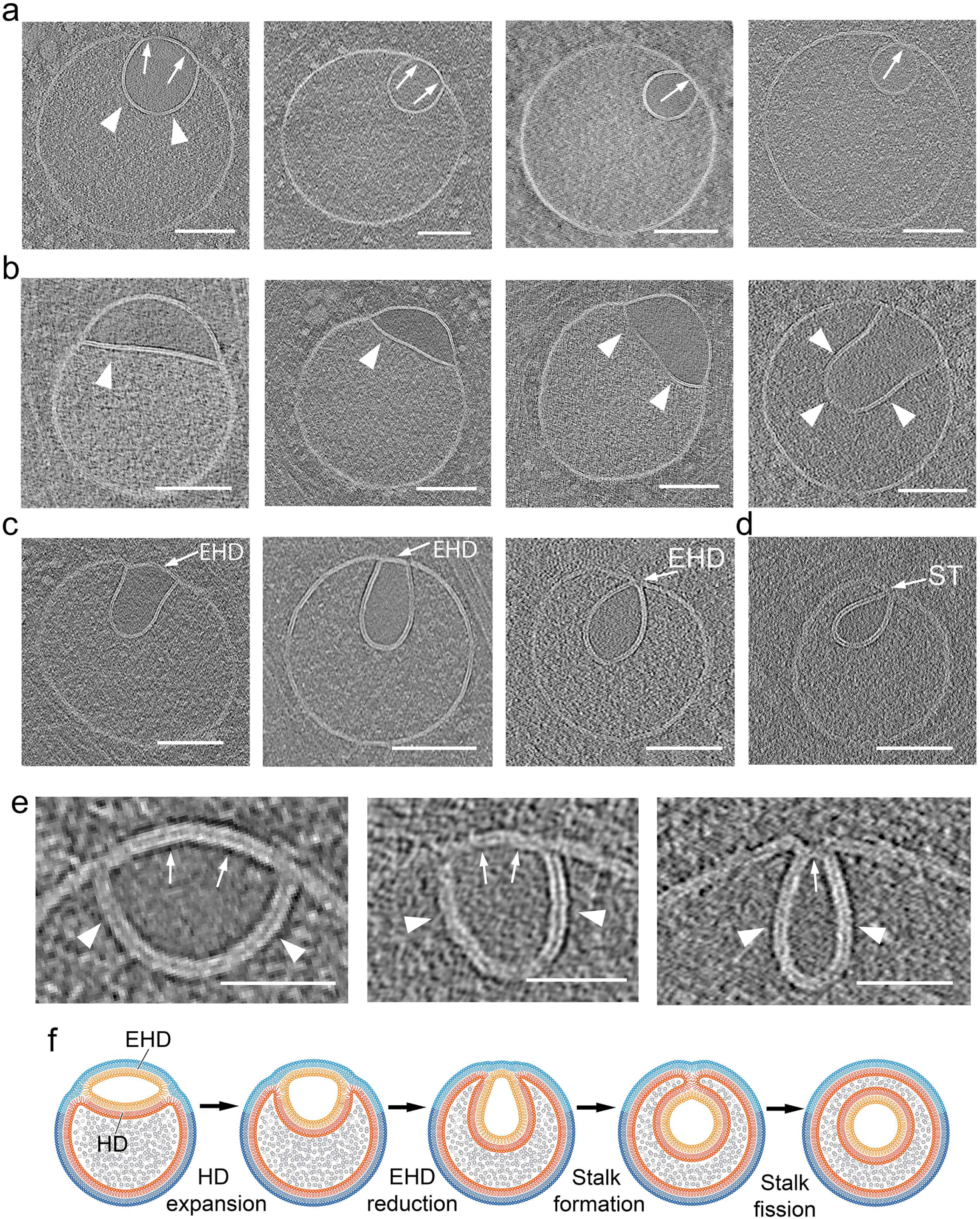
Range of morphological appearances of flipped hemifusomes and the proposed progression to form an intraluminal vesicle. Tomographic slices of various flipped hemifusomes (fHF) highlighting the variability in sizes and shapes across and within each hemifusome pair. a. Varied configurations of fHF showing various degrees of intraluminal budding of a vesicle while maintaining a round shape. b-d. Various degrees of curving and budding of the hemifusion diaphragm. The budding vesicle exhibits an elongated shape. In both scenarios, the intraluminal portion of the hemifused diaphragm expands, while the cytoplasmic side of the vesicle decreases in radius and forms an external segment of the bilayer shared by both vesicles. During this process, the cytoplasmic side of the lens-shaped structure transforms into an external hemifusion diaphragm (EHD), which reduces radially to form a stalk-like structure (ST). e. Close-up views of the curving and expansion of the hemifusion diaphragm (HD, arrowheads) and radial reduction of the EHD (arrows). f. Diagram depicting our proposed model, illustrating the progressive rounding and expansion of the hemifusion diaphragm (HD) and reduction of the EHD to form a stalk and ultimately scission to form an intraluminal vesicle. Scale bars: 200 nm.

Among the rounded hemifusomes, we observed a range of conformations (Fig. 4 and S4) that appear to correspond to various stages of HD expansion and intraluminal budding of the lens-shaped structure. Similarly, in flipped hemifusomes, we identified what we believe to be two modes of intraluminal budding. In the first (Fig. 4a), the smaller vesicle of the hemifusome pair is well-rounded, optimizing the volume-to-surface area ratio. In the second (Fig. 4b, c), the budding vesicle exhibits an elongated shape. We postulate that in both scenarios, the original hemifusome HD, with two exoplasmic leaflets, curves and expands to form the luminal hemifused vesicle, while the cytoplasmic portion of the bilayer shared by both vesicles decreases in radius (Fig. 4a-c) transforming into an external hemifusion diaphragm (EHD), which is ultimately comprised of both an exoplasmic and a cytoplasmic leaflet (Fig. 4f). The consistent presence of the EHD is a critical feature that distinguishes the luminal budding vesicles in the flipped hemifusome from the canonical ESCRT-based budding and formation of intraluminal vesicles ^5^. In the ESCRT model, the budding portion of the membrane forms an omega shape, with the cavity open to the cytoplasm (Fig. 2b) until the neck of the forming vesicle undergoes scission. Conversely, in the hemifusome, there is invariably an HD separating the cytoplasm from the lumen of the inwards-flipping vesicle (Fig. 4). Furthermore, in the ESCRT-based model, the content of the cavity corresponds to the portion of cytoplasm being captured (Fig. 2b), while in the reverse fusion process, the texture of the luminal content is smooth and lighter in contrast (Fig. 4), consistent with the “translucent” content of the smaller or lens-shaped vesicle of the hemifusome (asterisks in Fig. 3).

Intriguingly, the smaller EHDs appear to be less than double the thickness of the bilayer (Fig. 4c), akin to the reported dimensions of the canonical fission intermediate “stalk” structures ^14,20,21,34,35^. While we did not directly visualize the merging of the luminal membranes due to the resolution limitations of our tomograms, the point-like contact of the membranes as shown in Figure 4d is consistent in size to that of a stalk. This putative conversion of an HD into a stalk is akin to a reverse canonical fusion process ^18,20,36^ where the stalk precedes the formation of an HD. The diagram in Fig. 4f illustrates the range of conformations observed, and our proposed model showing how these conformations could fit within a progressive intraluminal budding of the translucent vesicle, with a concomitant reduction of the EHD until a stalk is formed. Like the scission of the intraluminal vesicle proposed in ESCRT-based intraluminal vesicle formation ^5^, it is plausible that the scission of the flipped hemifusome stalk then results in the pinching off of the vesicle and the formation of a free intraluminal vesicle.

### Exploring the relationship between hemifusomes and endosomes using gold nanoparticles

Given the similarity in size and content between the larger compartment of the hemifusome and similarly sized endosomal vesicles observed in our tomogram slices (Fig. S2a), we sought to explore the potential relationship between hemifusomes and endosomes by employing functionalized gold nanoparticles as tracers of endocytosis. Endosomes represent population of vesicles with remarkable variation in both their source and trajectories. They originate largely from the plasma membrane ^37,38^, but also form through the fusion of vesicles derived from other intracellular compartments, such as the trans-Golgi network ^27^. During receptor-mediated endocytosis, specific ligands bind to their corresponding receptors on the plasma membrane, and these receptor-ligand complexes are subsequently internalized into subsets of endosomes ^29^. This mechanism ensures that each endocytic pathway selectively internalizes specific cargo based on receptor-ligand interactions ^39^.

To investigate whether hemifusomes participate in the endocytic uptake pathway, we used three types of functionalized gold nanoparticles as tracers. These nanoparticles varied in size (5 nm and 15 nm) and surface functionalizations, including physiosorbed ferritin (Luna Nanotech), covalently bound ferritin, and a proprietary negatively charged polymer-coated nanogold (NanoPartz). After exposing the cells to the gold nanoparticles for various durations (ranging from 30 minutes to 3 hours), we observed their localization within coated pits, coated vesicles, endosomes, and lysosomes (Figs. 5 and S5). Strikingly, we were unable to detect gold nanoparticles within either vesicle of the hemifusomes (Fig. 5). Notably, we also observed many endosomes and lysosomes that did not contain gold (Fig. 5f), consistent with the diversity of endocytic pathways ^40–43^. Taken together, the results of our uptake experiment suggest that hemifusomes are not part of the transferrin endocytic pathway, but neither conclusively establish nor dismiss the possibility that hemifusomes represent a distinct subset of endosomes.

**Figure 5.**
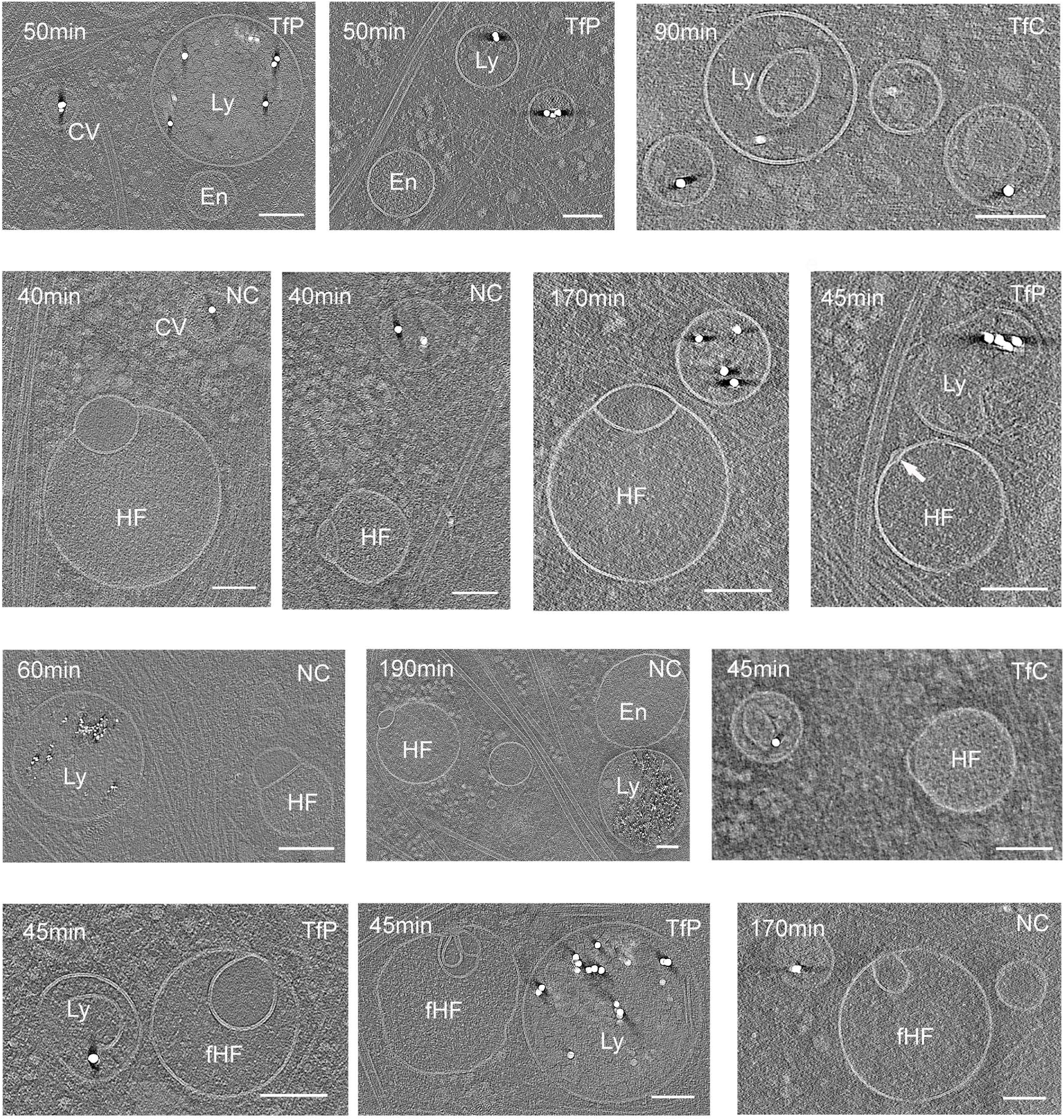
Hemifusomes are not part of the uptake and cargo transfer of endocytosed nanogold particles. Tomographic slices of pulse-chase experiments showing the distribution nanogold particles of various surface-functionalization and size in clathrin-coated pits or vesicles (CV), endosomes (En), and lysosomes (Lys), but absent from either vesicle compartment of the hemifusomes (HF) or flipped hemifusomes (fHF). T = time of incubation with the gold nanoparticles. TP = gold nanoparticles with transferrin physisorbed, TC = gold nanoparticles with transferrin covalently attached, and PC = gold nanoparticles with slightly positively charged non-reactive polymer. D = diameter of the gold nanoparticles. Scale bars: 200 nm.

### Proteolipid nanodroplet at the rim of the hemifusion diaphragm

As previously noted, tomographic analyses of hemifusomes reveal a single dense structure embedded within the hydrophobic interior of the bilayers at the three-way junction of the HD and the two heterotypic vesicles (Figs. 6a, 6b, 1b, 1c, S1c, 3a, and 3b). Detailed views show this dense structure, or nanodroplet, interfacing with the hydrophobic side of the exoplasmic leaflets of the hemifused vesicles and the HD (insets and close-up views in Figs. 6a and 6b), suggesting it contains hydrophobic components, likely lipids. The cytoplasmic surface and interior of the nanodroplet contain particulate structures (insets and close-ups in Figs. 6a and 6b), likely proteins, leading us to conclude that these nanodroplets are of mixed proteolipid composition.

**Figure 6.**
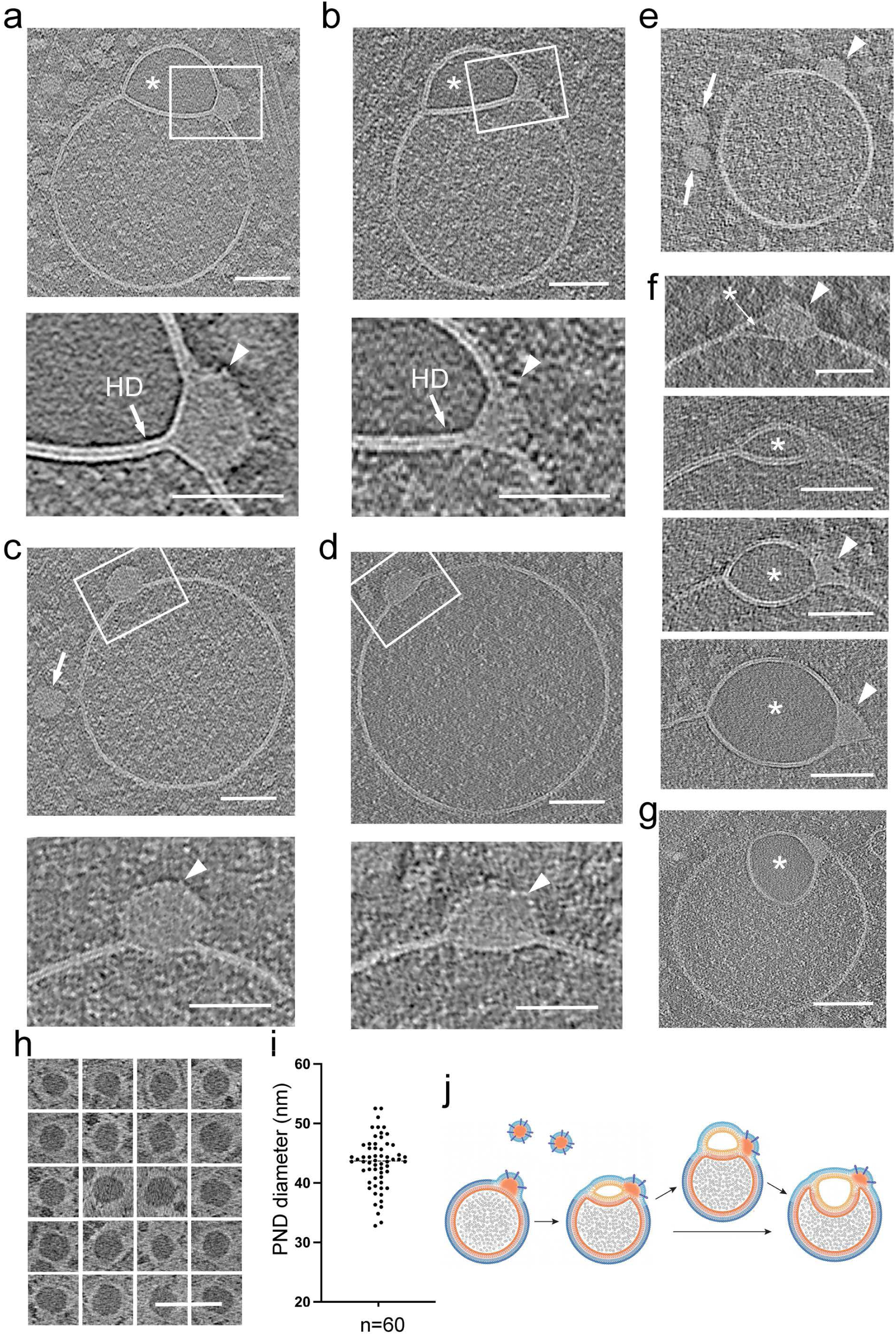
Proteo-lipid particles associated with hemifusomes are located at the rim of the hemifusion diaphragm. a and b. Tomographic slices and corresponding close-up views of hemifusomes (HF) with a prominent proteolipid nanodroplet (PND) at the rim of the hemifusion diaphragm (HD). Asterisk marks the smooth lumen of the smaller vesicle. Lower panels are close-up views of the area marked with a white rectangle showing the PND embedded in the hydrophobic core of the vesicle membrane and a series of particulate structures on its outer limit. These structural features reinforce the view that PNDs contain lipids and proteins. c and d. Tomographic slices and corresponding close-up views of PNDs fused to endosome-like vesicles. Arrows point to a PND free in the cytoplasm. Lower panels are high magnifications of the boxed regions, showing the PND encapsulated within the vesicle lipid bilayer. The outer leaflet enveloping the PND is decorated with protein particles (arrowheads). e. More examples of PNDs of uniform size and texture, within the cytoplasm (arrows) and fused to an endosome-like vesicle (arrowhead). f. Mid-cross-section of tomograms of a series of different hemifusomes showing a range of increasing size vesicles with a smooth lumen (asterisks) formed next to the PND site (arrowheads), suggesting that the smaller vesicle of the hemifusome may be forming by a de novo vesiculogenesis process. g. Tomogram slice of a flipped hemifusome showing the PND embedded at the three-way juncture of the hemifusion diaphragm and the membrane of the two vesicles. h. Panel of PNDs found free in the cytoplasm in the vicinity of endosomes and hemifusomes. Image contrast in this image is reversed to its original gray scale to show that the PNDs are electron dense or have a phase dark appearance. i. Plot of the diameter distribution of PNDs. The average diameter was calculated to be 42.4 ± 4.3 nm (n=60) j. Diagram illustrating the association of PNDs to endosomes to form a hemifusome and the progression to a flipped hemifusome.

Determining the frequency and distribution of nanodroplets around the entire HD is challenging due to the inherent missing wedge limitation (^44^ and Fig. S3i and j) of cryo-ET. Typically, the best tomograms cover about one-third of the hemifusome circumference, with the top and bottom thirds of the hemifusome missing from the tomogram (Fig. S3i and j). Consequently, even optimally oriented tomogram slices might miss these structures. Consistent with the estimation of one nanodroplet per hemifusome, a distinct nanodroplet was observed in about half of the ∼300 hemifusome tomograms examined, with no instances of two or more nanodroplets associated with a single HD.

Strikingly, similar nanodroplets are observed embedded within the hydrophobic interior of the bilayer of vesicles that are morphologically indistinguishable from endosomes (Figs. 6c–e). These lensed nanodroplets interface with the hydrophobic side of the exoplasmic leaflet of the vesicle bilayer, while the cytoplasmic boundaries are contiguous with the cytoplasmic leaflet of the vesicle bilayer (insets and close-ups in Figs. 6c–d).

We conducted a detailed examination of the diverse hemifusome and flipped hemifusome morphologies documented in our tomograms (Figs. 6f and 6g). Notably, we observed hemifusomes containing translucent vesicles, varying from minimal-sized pockets or cysts encapsulated within the bilayer adjacent to the nanodroplet, to progressively larger translucent hemifused vesicles in both direct hemifusome (indicated by asterisks in Fig. 6f) and flipped hemifusome configurations (asterisks in Fig. 6g). This observation raises the possibility that the insertion of the nanodroplet into endosomal or endosome-like vesicles might trigger the initiation, formation, or stabilization of a vesicle hemifused to the parent vesicle to form the hemifusomes. The images, which resemble a process of membrane blistering or vesiculogenesis, suggest a mechanism involving the import of water into the lumen, potentially explaining the translucent nature of this extended compartment within the hemifusome.

The diameter of these nanodroplets, measured in tomogram slices through their mid-portion, was calculated to be 42.4 ± 4.3 nm (n=60). Their size, electron density, and texture are comparable to similar particles observed in the surrounding cytoplasm (Figs. 6c and e, as well as Figs. 1b and c). Figure 6h shows a panel of representative cytoplasmic nanodroplets. The contrast in this figure is inverted to highlight that the content of nanodroplets is electron-dense or phase-dark compared to the surrounding cytoplasm and resembles that of lipid droplets (Fig. S5b), except that they are not limited by a lipid leaflet but rather a proteinaceous coat. Based on these features, which support the idea that the nanodroplets are proteolipidic in nature, and their nanoscale size, distinct from lipid droplets (Fig. 6i), we designate these particles as proteolipid nanodroplets (PNDs).

Our findings that the PND is associated with endosome-like vesicles, and with various configurations of hemifusome, lead us to postulate a model for PND-based vesiculogensis as a pathway for hemifusome biogenesis. In this model we hypothesize that PNDs, initially free in the cytoplasm, attach to endosomes or endosome-like vesicles. At this site, the PND contributes lipid and protein building blocks for the initiation and growth of the vesicle lensed within the bilayer, which ultimately grows to form the translucent vesicle characteristic of hemifusomes (Fig. 6j).

### Compound hemifusomes as hubs for the formation of complex multivesicular bodies

One-quarter of the total number of hemifusomes, both direct and flipped, identified showed one or more additional vesicles in a hemifused conformation with either or both vesicles of the initial hemifusome pair (88 direct, 48 flipped hemifusomes, and 42 compound out of 178 total hemifusomes; Fig. 7, S6 and S7). The occurrence of these compound hemifusomes further demonstrates the unexpected longevity of the HDs within the hemifusomes. We also observed several instances of hemifusomes with additional PNDs embedded in their membranes. Based on our hypothesized model of PNDs as hubs for vesicle biogenesis (Fig. 6), we propose that these additional PNDs are likely sites for the initiation of compound hemifusomes.

**Figure 7.**
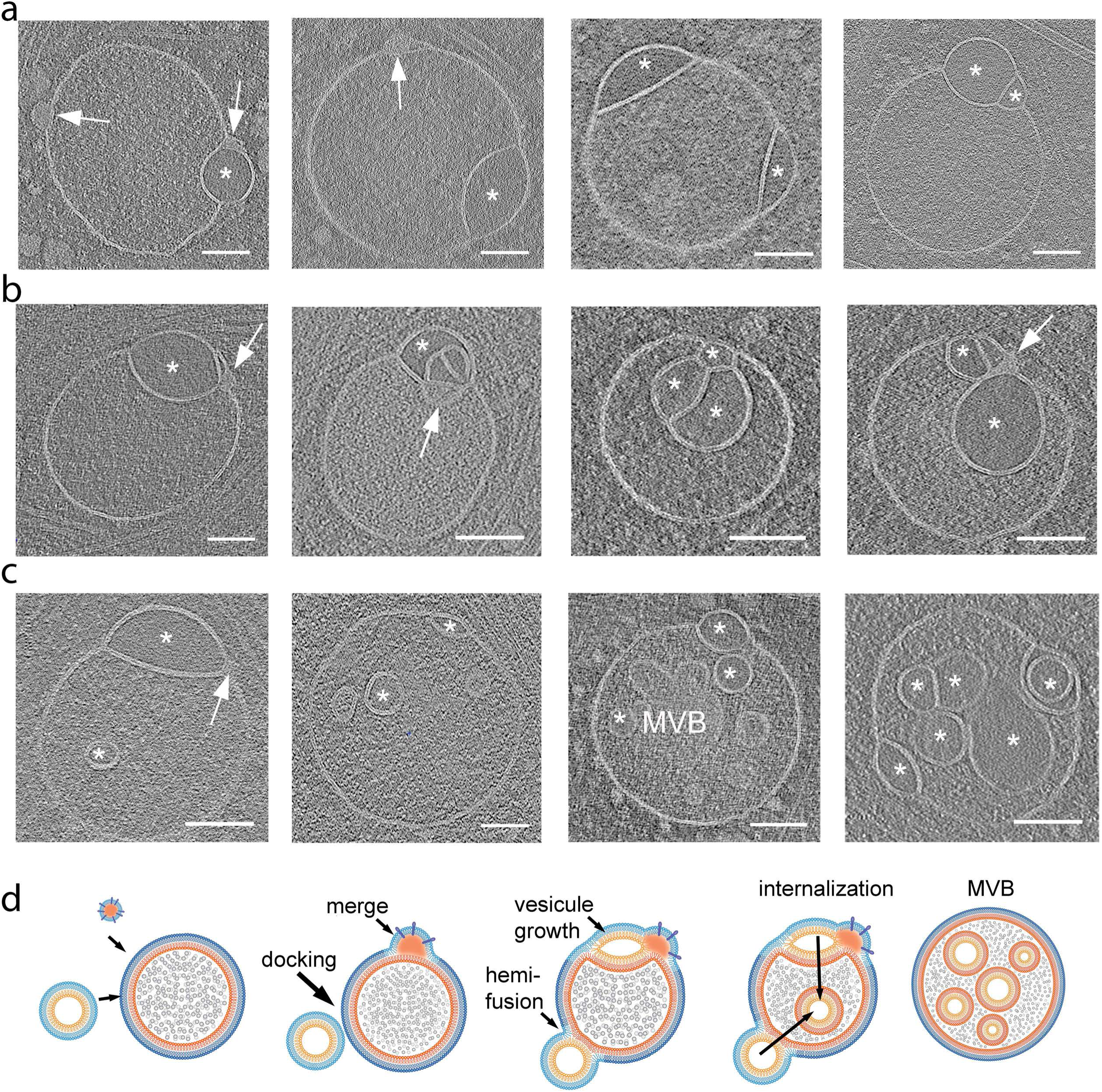
Compound hemifusomes as hubs for the formation of late endosomes and multivesicular bodies. a. Tomographic slices illustrating compound hemifusomes with one or more additional vesicles in a hemifused conformation with either of the two vesicles of the initial hemifusome pair. Asterisks mark vesicles with clear luminal content. Hemifusomes with additional PNDs embedded in their membranes (arrows) suggest these may act as a hub for the formation of compound hemifusomes. b. Compound hemifusion with flipped hemifusomes. Multiple hemifusion events coalesce to form very complex compound structures. Asterisks mark vesicles with clear luminal content. c. Hemifusomes comprising hemifused vesicles as well as multiple intraluminal vesicles. Asterisks mark vesicles with clear luminal content. Arrow points to PND. d. Diagram illustrating the proposed path for the formation of compound hemifusomes, followed by the flipping of the vesicle into the luminal side of the larger vesicle and subsequent scission, providing an alternative path to the formation of late endosomes and multivesicular bodies (MVBs). Asterisks mark vesicles with clear luminal content. Scale bar: 200 nm.

Multiple hemifusion events coalesce to form very complex compound structures, as shown in Fig. 7c. Some of these structures harbor a range of conformations of the direct and flipped hemifusomes, suggesting that the mechanism of addition of hemifused vesicles is stochastic in time and location within the hemifusome complex (Fig. 7c). We speculate that compound hemifusomes, followed by the flipping of the vesicle into the luminal side of the larger vesicle and subsequent scission, may provide an alternative path to the formation of intraluminal vesicles (Fig. 7c and d). One relevant observation is that the outer vesicle in the hemifusome often shows a subtle crenation (Fig S7c and d), suggestive of reduced turgor. This reduction in turgor would facilitate the inward budding of the lens-shaped vesicle postulated in our proposed model.

## Discussion

Through the application of cryo-ET to cultured mammalian cells, we have identified a previously unrecognized organelle complex with a unique membrane topology. This complex, which we have termed ‘hemifusome’, consists of hemifused vesicles sharing a large (∼160 nm) hemifusion diaphragm (HD). The presence of such large, long-lived HDs is particularly surprising, given the widely accepted view that HDs are small (less than 10 nm), unstable, and typically occur as transient intermediates during rapid vesicle fusion processes leading to content mixing ^20,21,34^. A second intriguing feature of hemifusomes is the consistent presence of the distinct 42 nm proteolipid nanodroplet (PND) embedded in the membrane of the hemifused vesicles at the rim of the HDs. This localization suggests a role for the PND in the formation, stabilization, or expansion of the hemifusion interface or in hemifusome biogenesis and dynamics.

The paired vesicles in each hemifusome are heterotypic in both size and luminal content. Hemifusomes present in two distinct conformations: a direct hemifusome, where the smaller vesicle is hemifused to the cytoplasmic side, and a flipped hemifusome, where the hemifused vesicle is internal and fused to the luminal side of the parent vesicle. Given their distinctive topology and variety of conformations, we predict that hemifusomes may play roles in protein and lipid sorting, recycling, and the formation of intraluminal vesicles.

One possibility for hemifusome formation is through the hemifusion of two pre-existing vesicles. This hypothesis is supported by the occasional observation of docked and hemifused endosomal and lysosomal vesicles with expanded HDs (Figs. 2c and 2d).

However, the smaller vesicles in the hemifusome consistently contain a translucent luminal content, distinct from the appearance of the luminal content in all other vesicular organelles (including endosomes and lysosomes). In fact, in all 308 tomograms, we did not observe similar individual translucent vesicles in the cytoplasm. The only vesicles we observed with similar translucent content were some of the intraluminal vesicles in multivesicular bodies (MVBs). The absence of free translucent vesicles in the cytoplasm or docked to endosomes challenges the hypothesis that hemifusome formation occurs through canonical SNARE-mediated vesicle fusion.

The range of hemifusome morphologies observed in our in situ cryo-ET data—spanning from slightly to highly deformed hemifused vesicles sharing an expanded HD—has not been previously described in situ. These morphologies align with stages of HD remodeling and expansion predicted by mathematical models, synthetic lipid systems, and in vitro observations of reconstituted systems ^18^, where it has been suggested that a growing HD experiences high tension and may rupture if subjected to osmotic pressure or high-curvature membrane stresses ^21,31,45^. Detailed in situ analyses of lipid systems indicate that the HD can stabilize into a “lens-shaped” complex, known as “dead-end hemifusion” ^18,21^, which encapsulates a vesicle within a bilayer. Our observation of such structures in cells confirms that this phenomenon occurs in biological membranes, and potentially serves a biological function beyond acting as an intermediate.

Based on our observations, we propose that in hemifusomes, the lens-shaped structures are long-lived and can be remodeled beyond the theoretical dead-end formulation, evolving into intraluminal vesicles. This process may involve the progressive intraluminal budding of the smaller vesicle through a mechanism distinct from the canonical ESCRT pathway. The presence of an extended HD, as opposed to the “omega“-shaped budding that captures a portion of the cytoplasm, as described in ESCRT-mediated intraluminal vesicle formation, highlights the unique nature of hemifusome-mediated vesicle internalization. Given the consistent presence of a PND free in the cytoplasm, attached to endosome-like vesicles, and at the rim of the hemifusome HD, we hypothesize that PND are hubs for hemifusome formation, acting as pockets of membrane building blocks that utilize endosomal vesicle bilayers as sites and templates to generate new hemifused vesicles (Fig. 6). We further argue that this PND-dependent process, which we term de novo “vesiculogenesis,” represents an alternative to canonical ESCRT-based vesicle budding and intraluminal vesicle formation (Fig. 7).

The ESCRT-model of intraluminal vesicle formation presents with several challenges ^5,9^, particularly regarding the formation of multivesicular bodies (MVBs) containing many intraluminal vesicles. Generating sufficient membrane area in these cases would require a large supply of lipids, which is unlikely to be sourced from a single vesicle through repeated ESCRT-based internal budding and scission without an additional lipid source. Moreover, while there is evidence supporting the role of ESCRT filaments in promoting membrane budding and scission, direct evidence explaining how ESCRTs facilitate the formation of numerous nascent budding vesicles remains elusive ^7^. The proposal of a PND-dependent de novo vesicle formation, although not yet validated, is an appealing alternative mechanism in addition to ESCRT-based vesicle internalization. It not only aligns with our observed results but also offers a potential explanation for how lipids can be added to generate complex MVBs. Future studies will focus further testing, verification and validation of this model using live-cell markers.

Regardless of their biogenesis—whether by hemifusion, de novo formation driven by PNDs, or another process—these novel vesicle complexes with two or more compartments separated by HDs expand the diverse repertoire of membrane remodeling processes mediated by the endolysosomal system. The HDs, with their unique bilayer formed by two exoplasmic leaflets, will impact the conformation and distribution of proteins according to their topological sensitivity^46,47^. They could be involved in various processes associated with endosome maturation ^48^, including lipid transfer ^49^ and lipid sorting ^50^, or could explain the existence of lipid raft-like microdomains that retain specific proteins ^43^. An intriguing possibility is the relationship between hemifusomes and recently reported lysosome-related organelles that contain an expansion compartment mediating zinc transporter delivery ^51^, although these organelles are micrometer-sized structures and appear much larger than hemifusomes. The heterotypic vesicles in hemifusomes may also be related to amphisomes ^42^.

In situ cryo-ET is arguably the most promising approach to obtaining native structural information. The overall quality of our images attests to the preservation of structure. For instance, our in situ images of clathrin-coated pits and vesicles (Figs. 1b, 2a, and S2a) directly reveal additional structural features beyond those visualized using the widely used approach of unroofing cells ^52,53^. In our in situ cryo-ET approach, we aimed to minimize artifacts or stress responses in cells that often result from sample handling and preparation. We achieved this by reducing sample manipulation to essential steps only. Specifically, the process involved a brief 2-3 second transfer of unperturbed cells from the tissue culture medium to the plunge-freezing apparatus, followed by a 4-6 second blotting phase in the humid chamber of the apparatus prior to vitrification. It is thus unlikely that hemifusomes and their associated PNDs are artefacts. However, if hemifusomes are indeed formed because of this minimal handling, they can be regarded as part of a novel, rapid cellular stress response that must be considered in the evaluation of in situ cryoET data. It is more likely that hemifusomes and their HDs were previously overlooked in conventional electron microscopy due to fixation and dehydration steps, which may alter their stability, longevity, or appearance.

Our cryo-ET observations have been limited to the thin regions at the cellular periphery. Future research should focus on determining whether hemifusomes and compound hemifusomes are present in other cellular regions and on elucidating the molecular mechanisms underlying their formation, stability, and function, as well as their broader implications for cellular physiology and pathology.

## Supplementary Figure Legends

**Supplementary Figure 1.**
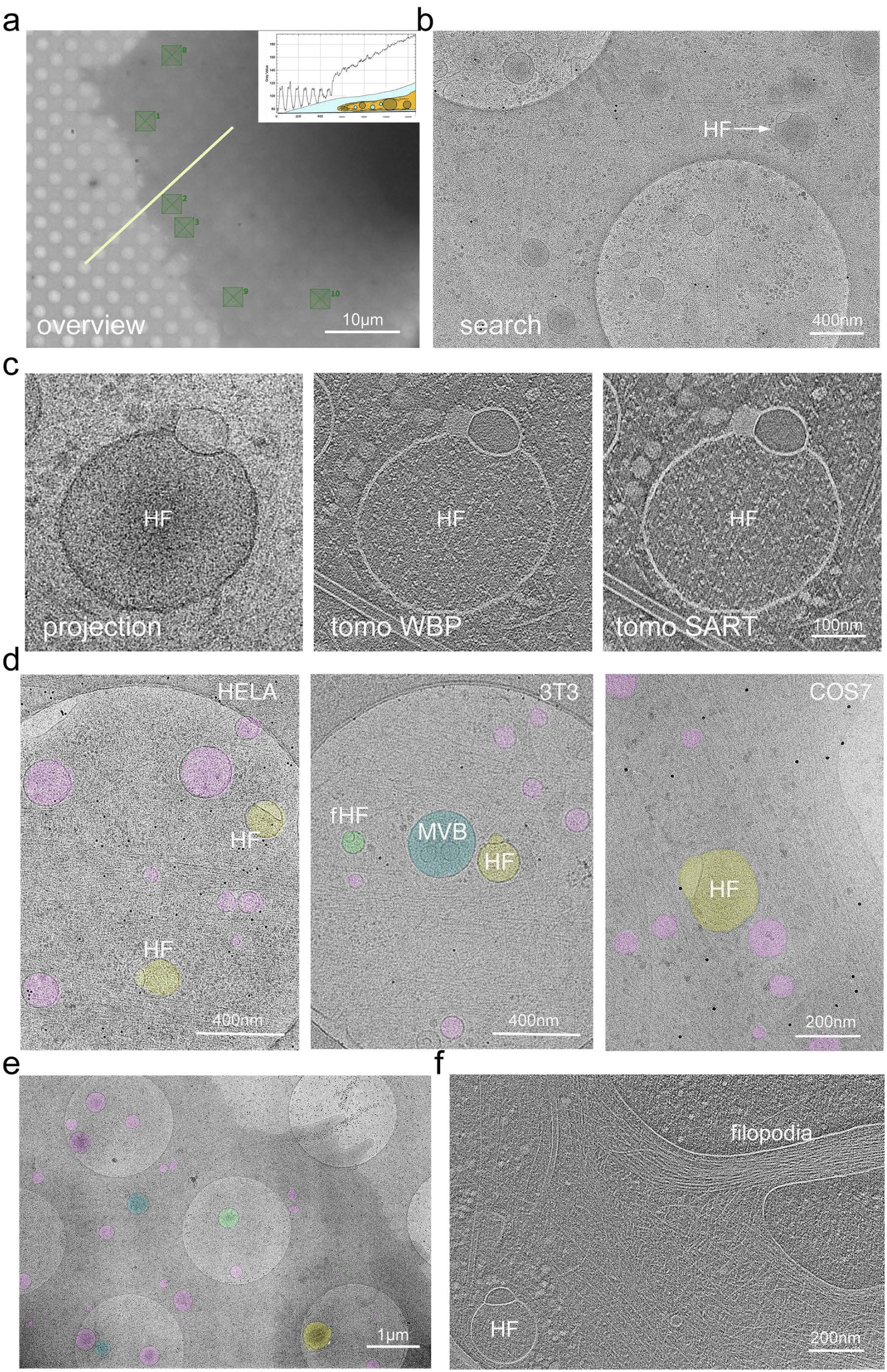
Workflow of image acquisition and distribution of hemifusomes and in various cell lines. a. Overview cryo-EM image of the leading edge of a COS7 cell on a CryoEM grid showing the gradient of ice thickness (inset) and examples of the targets selected for acquisition of “search” images and tilt series for tomography. Scale bar: 10 μm. b. View of a representative “search” image. Hemifusome labeled HF. Scale bar: 400 nm. c. Close-up view of a representative “projection” image from a raw tilt series acquisition and corresponding tomographic reconstruction using either the weighted back projection (WBP) or simultaneous algebraic reconstruction technique (SART). Scale bar: 100 nm. d. Representative search views of HELA 3T3, and COS7 cells with vesicular organelles highlighted in color: endosomes (EN, in pink); multivesicular bodies (MVB, in blue); hemifusomes (HF, in yellow); and flipped hemifusomes (fHF, in green). The COS7 cell image in this panel was used to target and collect the tilt series, and reconstruct the tomogram shown in Figure 1b. Scale bar: 400 nm and 200 nm. e. Representative, low magnification, overview of a COS7 cell periphery illustrating the frequency of hemifusomes compared to the other vesicular organelles in the cell border. Scale bar: 1 μm. f. Tomogram slice of a COS7 cell leading edge showing a hemifusome (HF) in a region of cytoplasm with a filopodia and a dense cytoskeletal network. Scale bar: 200 nm.

**Supplementary Figure 2.**
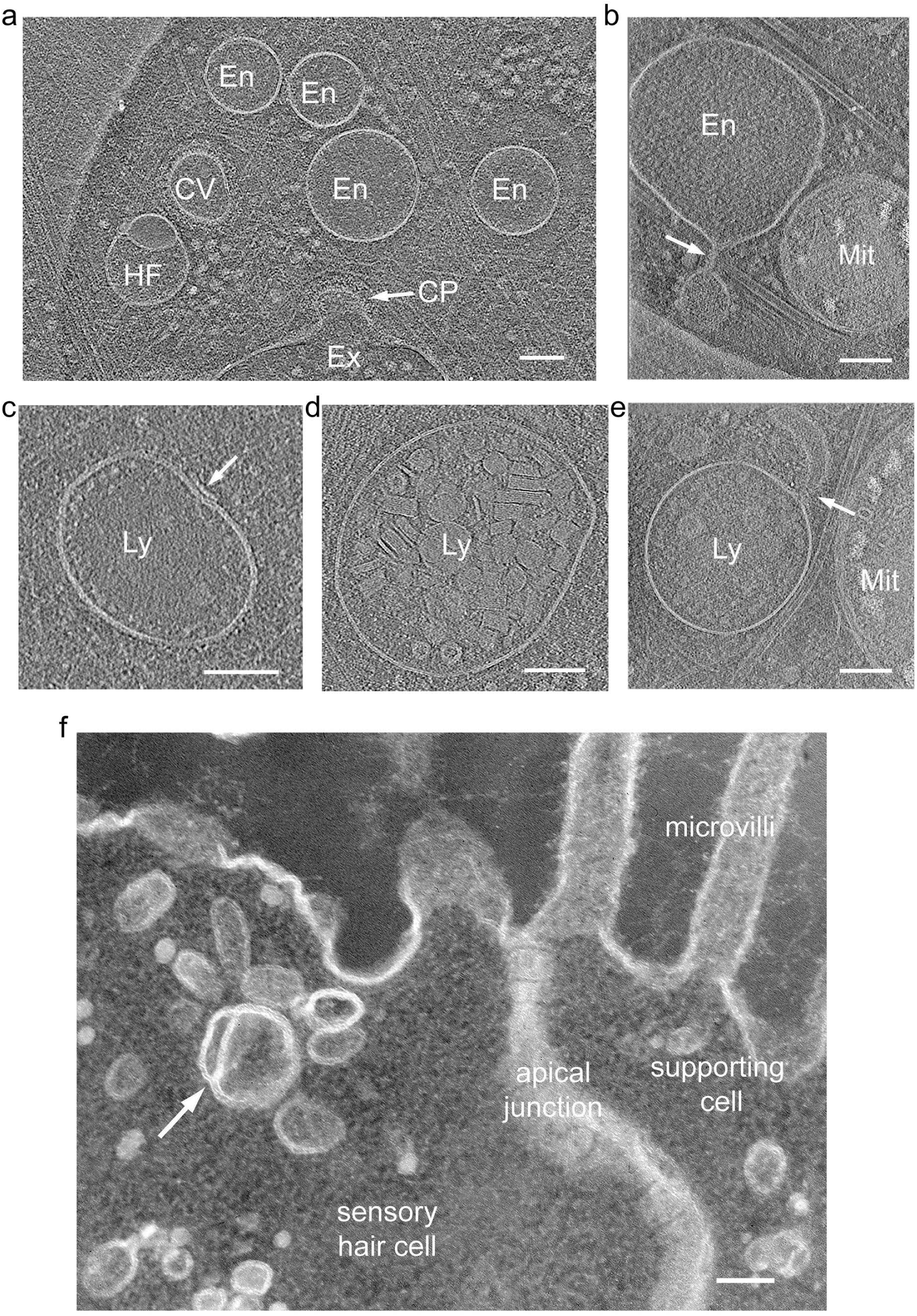
Hemifusome among endolysosomal vesicles in a cultured cell line and in a native epithelial cell. a-e. Cryo-ET images of a COS-7 cell showing organelles at the cell periphery, including: a hemifusome (HF) near a clathrin-coated pit (CP), a clathrin-coated vesicle (CV), and variably sized endosomes (a); Endosome (En) where a vesicle appears to be pinching off as shown by arrow (b); Lysosome (Ly) with slight membrane invagination decorated with protein particles (c); Lysosome (Ly) with complex content; Lysosome with tubulation depicted by arrow (d). b. Conventional electron microscopy of a thin section of fixed and plastic-embedded epithelial cell from the frog macula. The image shows the apical cytoplasm of an epithelial cell with a hemifused pair of vesicles among several endosomal vesicles. Scale bars: 100 nm.

**Supplementary Figure 3.**
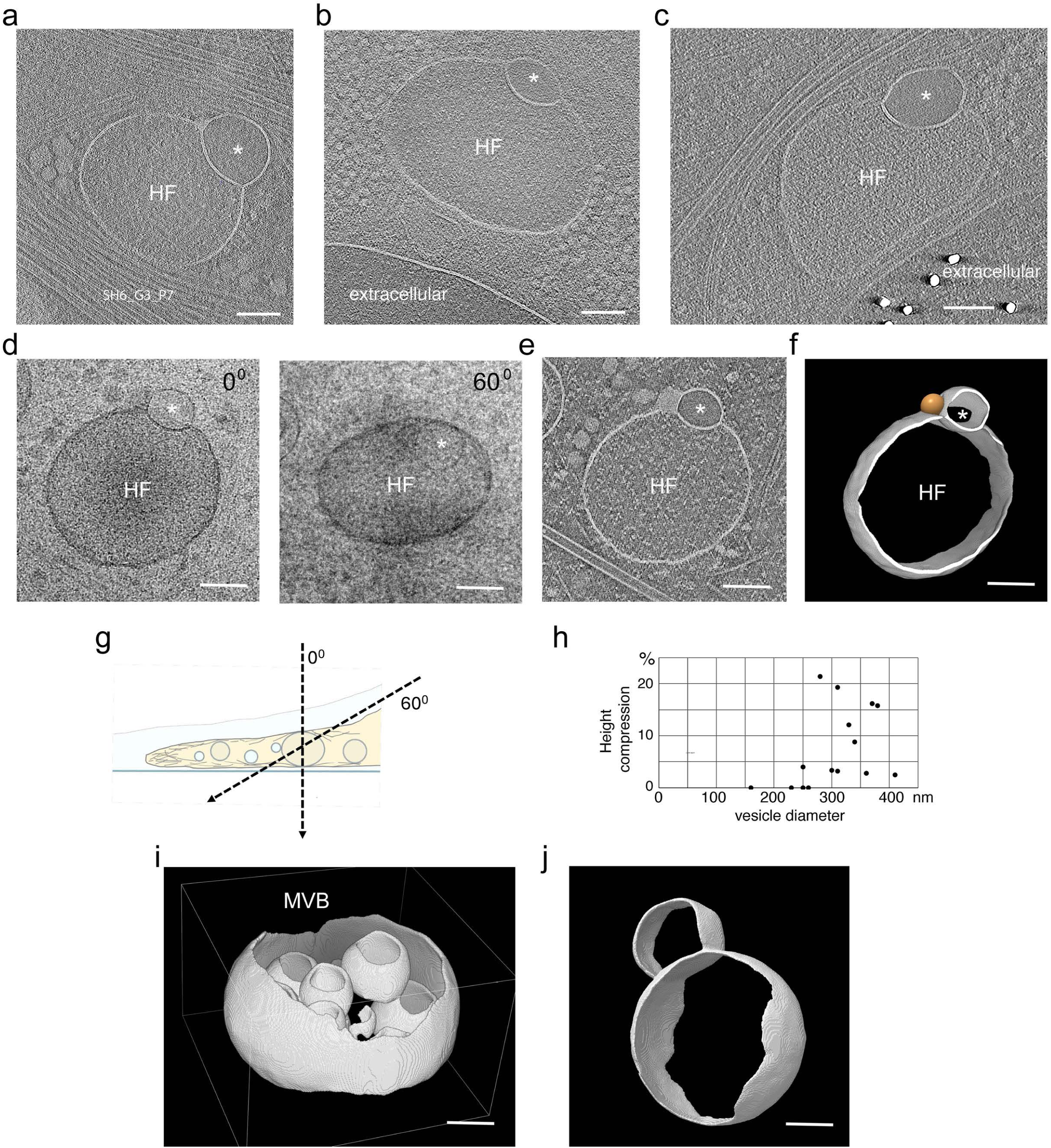
Morphological variability and deformability of hemifusomes. a-c. Series of cryo-ET mid-cross-sections showing hemifusomes laterally compressed by cytoskeletal elements (a), the plasma membrane (b), or both (c). Asterisks mark vesicles with clear luminal content. d. Single projection images from a raw tilt series at 0° and at 60° relative to the cryo-EM grid as illustrated in g showing the vertical compression of the hemifusome. e. Reconstructed tomogram showing slight compression of the vesicle. f. 3D-rendered segmentation of hemifusome in (e). The golden sphere in represents the proteolipid particle commonly located at the rim of the hemifusion diaphragm. g. Schematic illustrating the missing wedge effect limitations of cryo-ET, where membranes cannot be visualized above and below a narrow strip of the total vesicle volume. Scale bars: 100 nm h. Plot of compression factor i-j. 3D views of segmented volumes of a multivesicular body (i) and a hemifused pair of vesicles (j) illustrating the missing wedge or missing cone effect in the reconstructed tomograms.

**Supplementary Figure 4.**
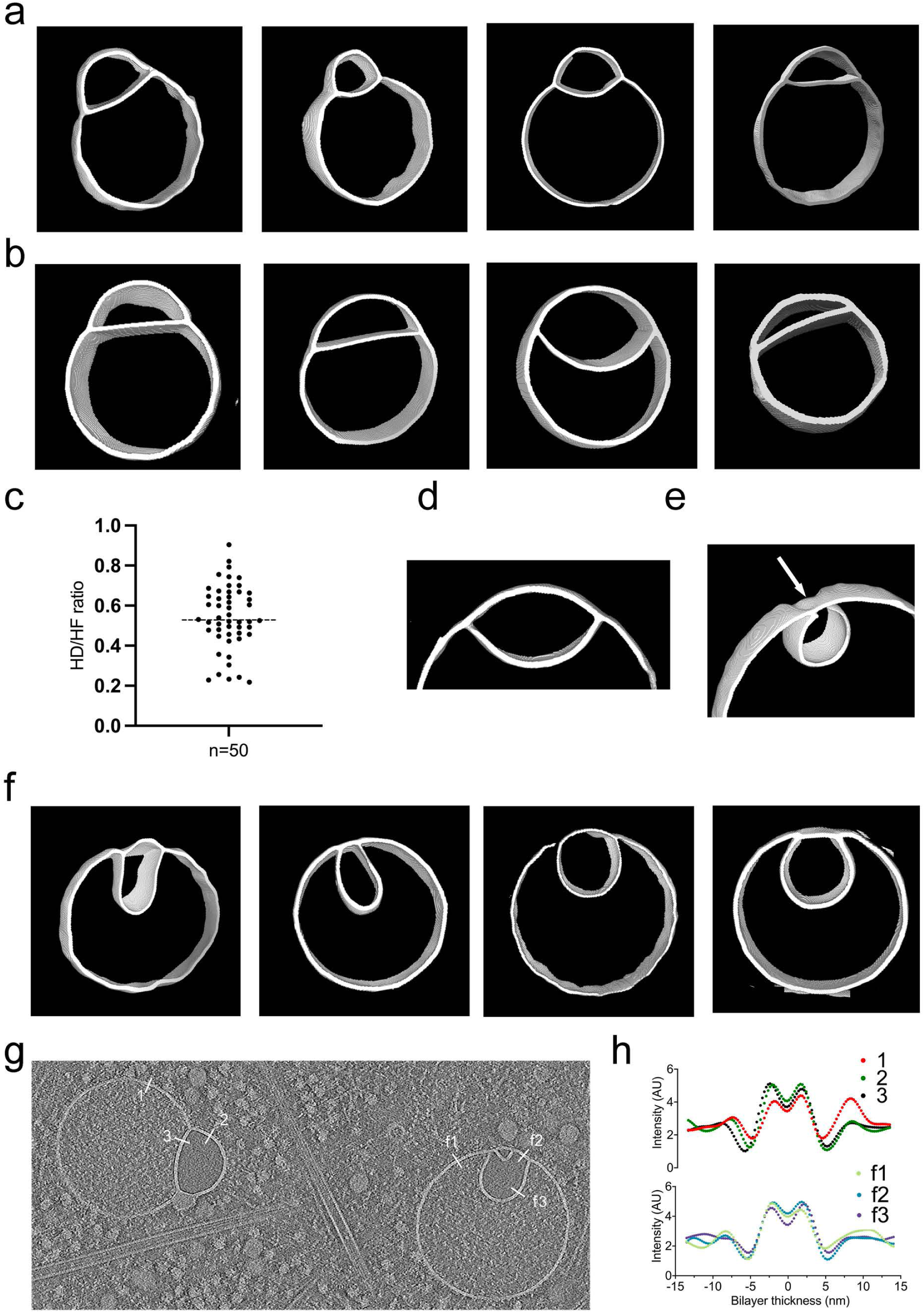
Range of morphologies of hemifusomes and flipped hemifusomes highlighted in segmented tomograms. a-b. 3D views of segmented hemifusome membranes illustrating various size ratios, roundness, and curvature of the hemifusion diaphragm. These reconstructions are from hemifusomes shown in the main figures. c. The ratio of diameters of the hemifusion diaphragm (HD) and larger vesicle of the hemifusome (HF). n = 50 hemifusomes. d. Close-up view of the lens-shaped structure of the smaller vesicle of the hemifusome. e. 3D views of segmented flipped hemifusome membranes illustrating various degrees of inward budding of the smaller vesicle into the lumen of the larger vesicle. f. Close-up view of the external hemifusion diaphragm (arrow). Note that some membranes of the flipped hemifusomes show subtle crenation indicative of reduced turgor. g. Image of direct and flipped hemifusomes to illustrate where measurements were made for membrane bilayer thickness (lines drawn). h. Graphs depicting pixel intensity along the lines drawn in (g). The maxima in each trace represent each leaflet of the bilayer. Bilayer thicknesses measured: 1= 4.149 nm, 2= 4.105 nm, 3= 4.546 nm, f1= 4.135, f2= 4.142, f3= 4.55.

**Supplementary Figure 5.**
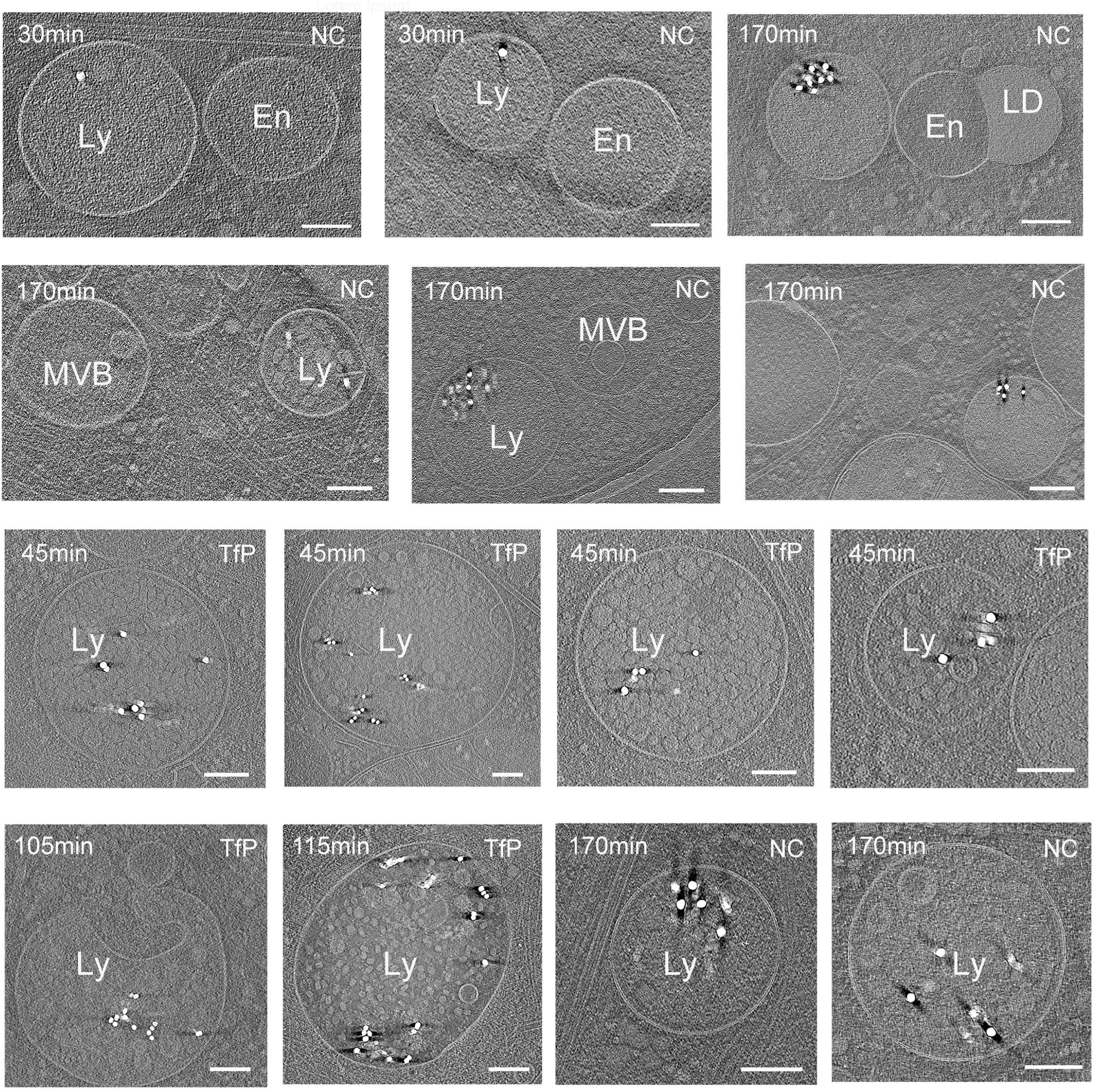
Pulse-chase of nanogold particles in the endocytic pathways. Tomographic slices examples of pulse-chase experiments showing the uptake of nanogold particles of various surface functionalization and sizes (small particles = 5nm and large particles = 15 nm) in endosomes (En), and lysosomes (Ly), but absent from MVBs and a vesicle hemifused to a lipid droplet. Gold particles are enriched in mature, large lysosomes. T = time of incubation with the gold nanoparticles. TfP = gold nanoparticles with transferrin physiosorbed and NC = gold nanoparticles with slightly negatively charged non-reactive polymer. Scale bars: 200 nm.

**Supplementary Figure 6.**
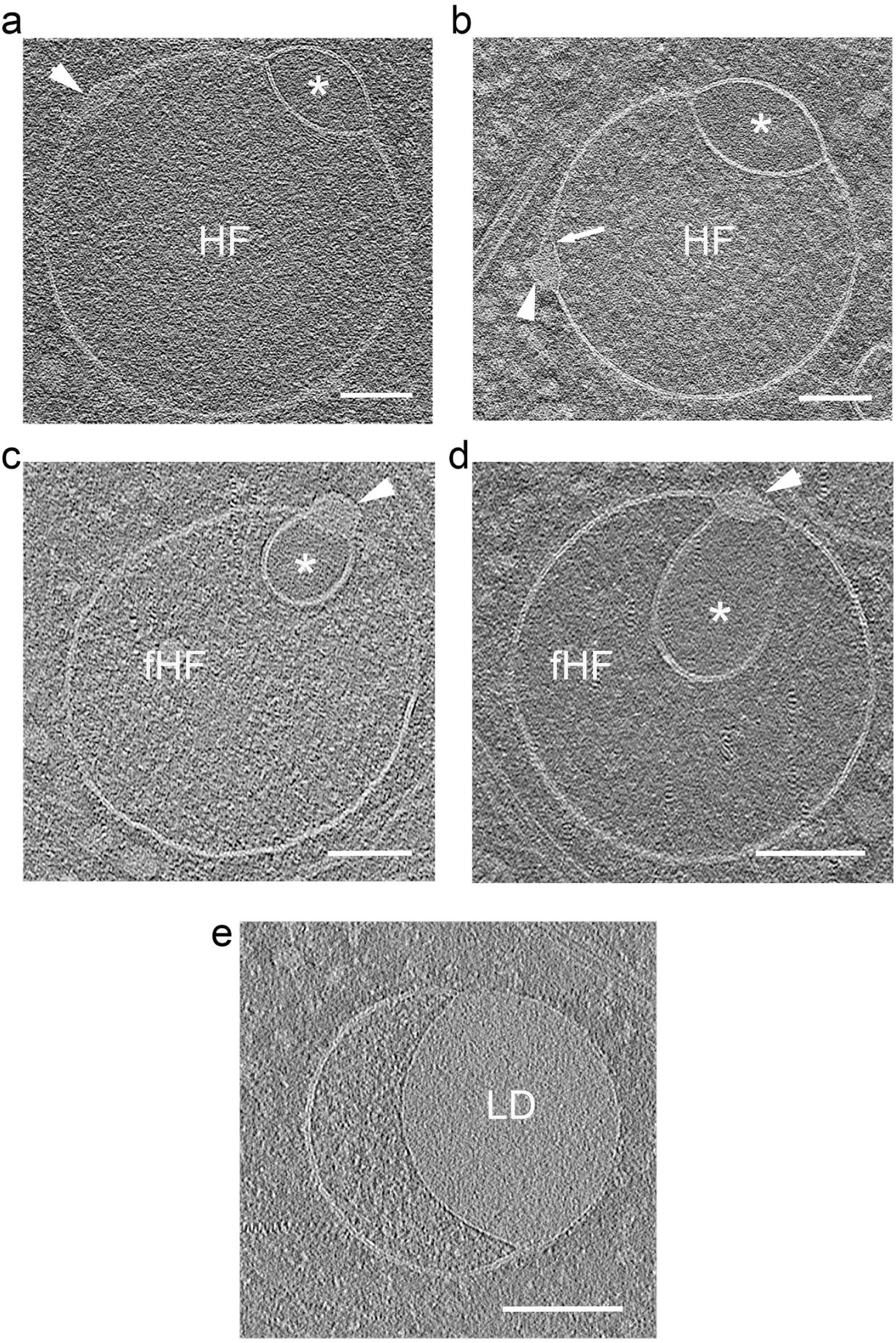
Proteolipid particles embedded in the membrane of hemifusomes, and flipped hemifusomes. a-b. Tomographic slice representative of additional PNDs embedded in the membrane of hemifusomes (HF) at sites away from the initial HD. c-d. Tomographic slice representatives of PND at the site of contact of the inner and outer vesicles of a flipped hemifusome (fHF). Asterisks mark vesicles with clear luminal content. e. Hemifused endosome and lipid droplet (LD). The texture and contrast of the LD is very comparable to that of the PNDs in (a-d), highlighting that the PND very likely is comprised of lipids, but with a proteinaceous coat that contrasts with the lipid monolayer surrounding the LD.

**Supplementary Figure 7.**
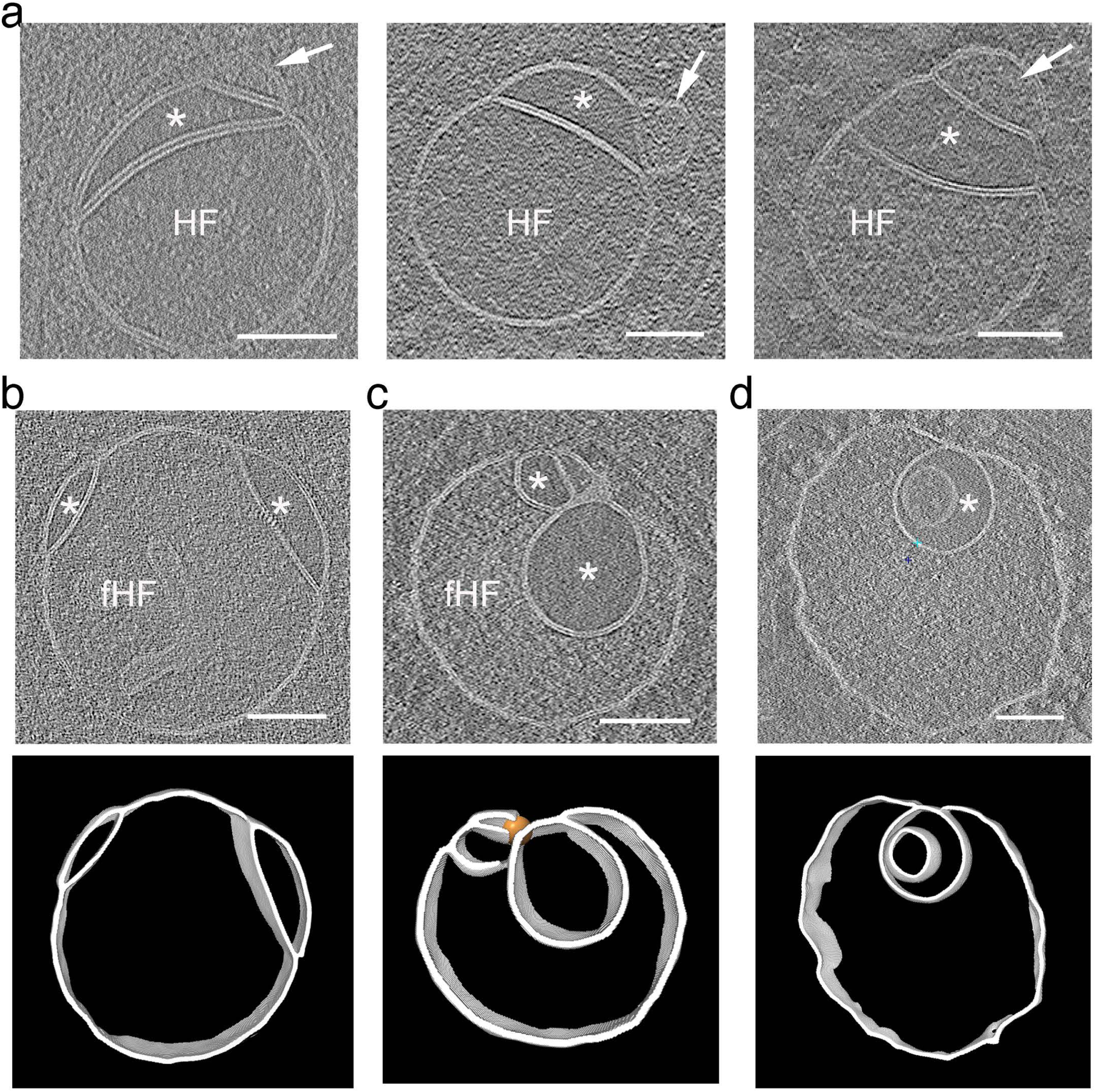
Additional examples of compound hemifusomes. a. Panel of cryo-ET slices showing compound hemifusomes and flipped hemifusomes, consistent with the hypothesis that they evolve into complex multivesicular bodies (MVB). Asterisks mark vesicles with clear luminal content. b - d. Panel of cryo-ET images (top) and corresponding segmentation (bottom) of compound hemifusomes (HF), (b), compound flipped hemifusomes (fHF), (c), and compound fHF with free intraluminal vesicles (d). Often, the compound flipped hemifusome shows a slightly crenated membrane indicative of reduced turgor pressure. Scale bars: 200 nm.

## Materials and Methods

### Cell Culture and Cryo-Preparation

Four cell lines were used: COS-7 (ATCC CRL-1651), HeLa (ATCC CCL-2), NIH/3T3 (ATCC CRL-1658), and Rat1 cells stably expressing claudin2-GFP as previously described ^54,55^. Cells were cultured in Dulbecco’s modified Eagle’s medium (DMEM) supplemented with GlutaMAX (Thermo Fisher Scientific, 10569010) and 10% heat-inactivated fetal bovine serum (HI-FBS; Thermo Fisher Scientific, 10082147) at 37°C in a 5% CO₂ humidified incubator. Cells were detached from culture flasks (Falcon, 353109) by rinsing with Dulbecco’s phosphate-buffered saline (DPBS; Thermo Fisher Scientific, 14190144) and treating with 0.05% Trypsin-EDTA (Thermo Fisher Scientific, 25300054) at 37°C for 2-3 minutes.

For cryo-preparation, gold EM grids with Quantifoil holey carbon film (Quantifoil, R2/1, 300 mesh, gold; Electron Microscopy Sciences, Q3100AR1; or Quantifoil, R3.5/1, 200 mesh, gold; TED PELLA, 660-200-AU-100) were mounted onto PDMS stencil grid holders (Alveole). PDMS stencils attached to coverslips were sterilized under a UV lamp for 2 hours. EM grids and stencils were glow-discharged (PELCO easiGlow, TED PELLA) for 30 seconds at 15 mA and coated with 10 μg/mL fibronectin (Fisher Scientific, 3416351MG) in DPBS for 2 hours at room temperature. Cells were seeded on 35 mm tissue culture dishes (Falcon, 353001) containing 2-3 fibronectin-coated EM grids at a density of 100,000 cells per dish for COS-7, NIH/3T3, and Rat1 cells, and 80,000 cells per dish for HeLa cells. After overnight incubation at 37°C and 5% CO₂, the medium was replaced with serum-free DMEM before plunger freezing.

Grids were carefully picked up using snap-lock forceps (Leica Microsystems, 16706435) and blotted with Whatman grade 40 filter paper (Whatman, 1440150). A 1 μL solution of 10 nm colloidal gold coated with BSA in DMEM was added to the grids ^56^. Samples were back-blotted for 6 seconds with Whatman grade 1 filter paper (Leica Microsystems, 16706440) and plunge-frozen into liquid ethane using a Leica EM GP plunger (Leica Microsystems) at 25°C, 85% humidity, and −176°C. Samples were stored under liquid nitrogen until data collection.

### Nanogold Tracing Experiments

Gold nanoparticles used in uptake studies included cell uptake, slightly negatively charged, polymer-functionalized gold nanoparticles (NanoPartz, CO), and transferrin-functionalized gold nanoparticles (Luna Nanotech, ON) as listed below. COS-7 cells were transfected with 1 μg of transferrin receptor mCherry-TFR-20 (Addgene, 55144) using Lipofectamine LTX (Thermo Fisher Scientific, 15338100) overnight. Functionalized nanoparticles were sonicated for 5 minutes before use and diluted in fresh DMEM with 10% HI-FBS for uptake experiments. Details of gold nanoparticle concentrations and uptake times are as follows:

- 15 nm Human transferrin covalently functionalized gold nanoparticles with AFDye 647 (GNP-TF-15-H-1-AF6): 1:10 dilution, uptake time: 30 min to 1 hr 15 min
- 15 nm Human transferrin physisorbed gold nanoparticles with AFDye 647 (GNP-TF(PA)-15-H-1-AF6): 1:10 dilution, uptake time: 45 min to 1 hr 45 min
- 5 nm cell uptake polymer-functionalized gold nanoparticles with AFDye 488 (PCU11-5-AF488-NCU-PBS-50-1-CS): 1:50 or 1:200 dilution, uptake time: 1 hr to 3 hr 15 min
- 15 nm cell uptake polymer-functionalized gold nanoparticles with AFDye 488 (PCU11-15-AF488-NCU-PBS-50-1-CS): 1:150 or 1:100 dilution, uptake time: 30 min to 5 hrs

### Cryo-ET Imaging

Vitrified samples were imaged using a 300 keV Titan Krios transmission electron microscope (Thermo Fisher Scientific) equipped with a Bioquantum post-column energy filter (Gatan) operated in zero-loss mode with a slit width of 20 eV and a defocus range of 2.5-4 μm.. Tomo control software (Thermo Fisher Scientific) was used to record dose-symmetric tilt series ^57^ from −60° to +60° at either 3° or 2.5° increments. Tilt series images were collected using a K3 Summit direct electron detector (Gatan) at ×27,000 and ×31,000 magnifications under low-dose conditions in counting mode (6 or 10 frames per tilt series image, 0.05 s per frame, 2.16 or 2.69 Å pixel size). The cumulative electron dose per tilt series was limited to under 120 e-/Å².

### Data Processing

K3 movie frames were dose-weighted and motion-corrected using Motioncorr2, then merged using the IMOD software package ^58^ to generate the final tilt series data. Frames affected by drift or blockage were excluded from reconstruction. Preprocessing and 3D reconstruction were performed using IMOD or AreTomo ^59^. Tilt series images for hemifusion reconstructions were aligned using 10 nm gold nanoparticles as fiducials with IMOD or AreTomo2 for automated marker-free alignment. 3D reconstructions were calculated using weighted back-projection (WBP) algorithm from IMOD or AreTomo2, and simultaneous algebraic reconstruction technique (SART) algorithm from AreTomo2 to enhance the contrast. Tomograms were selected based on their effectiveness in accentuating hemifusion characteristics. For visualization of raw tomographic slices, tomograms were denoised using weighted median filter (smooth filter) implemented in IMOD ^58^ to enhance the contrast for WBP reconstructed tomograms. For the visualization and analysis of cellular membranes, we used MemBrain v2 package ^60^. MemBrain-seg module’s automated segmentation and Surface-Dice loss function enhanced the membrane connectivity on the Cryo-ET data ^60^. Isosurface renderings and curation of the segmentation were performed using UCSF ChimeraX ^61^ and Amira ^62^.

## Author contributions

The project was conceptualized by BK and SE. Experimental work was performed by SH and AT. Cryo-electron microscopy was done by BK. Tomograms were reconstructed by AT. The manuscript was written by BK and SE. All authors assisted in editing the manuscript and contributed to data analysis.

## Acknowledgment

We thank Dr. Michael Purdy and the UVA Molecular Microscopy facility for expert assistance and the use of the Krios electron microscope. This work utilized the computational resources of the NIH HPC Biowulf cluster (http://hpc.nih.gov). This project was supported by NIDCD-NIH-IRP funds Z01-DC000002 to BK, SH, and ATT and by and the Center for Cell and Membrane Physiology, School of Medicine, at the University of Virginia through a start-up grant to S.E

## Notes

### Competing Interest Statement

The authors have declared no competing interest.

## References

1 Knorr, R. L., Mizushima, N. & Dimova, R. Fusion and scission of membranes: Ubiquitous topological transformations in cells. Traffic 18, 758–761 (2017). 10.1111/tra.12509

2 Saffi, G. T. & Botelho, R. J. Lysosome Fission: Planning for an Exit. Trends Cell Biol 29, 635–646 (2019). 10.1016/j.tcb.2019.05.003

3 Knorr, R. L., Lipowsky, R. & Dimova, R. Autophagosome closure requires membrane scission. Autophagy 11, 2134–2137 (2015). 10.1080/15548627.2015.1091552

4 Piper, R. C. & Katzmann, D. J. Biogenesis and function of multivesicular bodies. Annu Rev Cell Dev Biol 23, 519–547 (2007). 10.1146/annurev.cellbio.23.090506.123319

5 Olmos, Y. The ESCRT Machinery: Remodeling, Repairing, and Sealing Membranes. Membranes (Basel) 12 (2022). 10.3390/membranes12060633

6 Jahn, R., Cafiso, D. C. & Tamm, L. K. Mechanisms of SNARE proteins in membrane fusion. Nat Rev Mol Cell Biol 25, 101–118 (2024). 10.1038/s41580-023-00668-x

7 Frankel, E. B. & Audhya, A. ESCRT-dependent cargo sorting at multivesicular endosomes. Semin Cell Dev Biol 74, 4–10 (2018). 10.1016/j.semcdb.2017.08.020

8 Hernandez, J. M. et al. Membrane fusion intermediates via directional and full assembly of the SNARE complex. Science 336, 1581–1584 (2012). 10.1126/science.1221976

9 Marsh, M. & van Meer, G. Cell biology. No ESCRTs for exosomes. Science 319, 1191–1192 (2008). 10.1126/science.1155750

10 Banushi, B. & Simpson, F. Overlapping Machinery in Lysosome-Related Organelle Trafficking: A Lesson from Rare Multisystem Disorders. Cells 11 (2022). 10.3390/cells11223702

11 Klein, S. et al. IFITM3 blocks influenza virus entry by sorting lipids and stabilizing hemifusion. Cell Host Microbe 31, 616–633 e620 (2023). 10.1016/j.chom.2023.03.005

12 Gruenberg, J. Life in the lumen: The multivesicular endosome. Traffic 21, 76–93 (2020). 10.1111/tra.12715

13 Carter, S. D. et al. Ribosome-associated vesicles: A dynamic subcompartment of the endoplasmic reticulum in secretory cells. Sci Adv 6, eaay9572 (2020). 10.1126/sciadv.aay9572

14 Chlanda, P. et al. The hemifusion structure induced by influenza virus haemagglutinin is determined by physical properties of the target membranes. Nat Microbiol 1, 16050 (2016). 10.1038/nmicrobiol.2016.50

15 Cornell, C. E., Mileant, A., Thakkar, N., Lee, K. K. & Keller, S. L. Direct imaging of liquid domains in membranes by cryo-electron tomography. Proc Natl Acad Sci U S A 117, 19713–19719 (2020). 10.1073/pnas.2002245117

16 Mageswaran, S. K., Yang, W. Y., Chakrabarty, Y., Oikonomou, C. M. & Jensen, G. J. A cryo-electron tomography workflow reveals protrusion-mediated shedding on injured plasma membrane. Sci Adv 7 (2021). 10.1126/sciadv.abc6345

17 Martin-Solana, E. et al. Ribosome-Associated Vesicles promote activity-dependent local translation. bioRxiv (2024). 10.1101/2024.06.07.598007

18 Warner, J. M., An, D., Stratton, B. S. & O’Shaughnessy, B. A hemifused complex is the hub in a network of pathways to membrane fusion. Biophys J 122, 374–385 (2023). 10.1016/j.bpj.2022.12.003

19 Tsai, H. H., Chang, C. M. & Lee, J. B. Multi-step formation of a hemifusion diaphragm for vesicle fusion revealed by all-atom molecular dynamics simulations. Biochim Biophys Acta 1838, 1529–1535 (2014). 10.1016/j.bbamem.2014.01.018

20 Joardar, A., Pattnaik, G. P. & Chakraborty, H. Mechanism of Membrane Fusion: Interplay of Lipid and Peptide. J Membr Biol 255, 211–224 (2022). 10.1007/s00232-022-00233-1

21 Golani, G. & Schwarz, U. S. High curvature promotes fusion of lipid membranes: Predictions from continuum elastic theory. Biophys J 122, 1868–1882 (2023). 10.1016/j.bpj.2023.04.018

22 D’Agostino, M., Risselada, H. J., Lurick, A., Ungermann, C. & Mayer, A. A tethering complex drives the terminal stage of SNARE-dependent membrane fusion. Nature 551, 634–638 (2017). 10.1038/nature24469

23 Morandi, M. I. et al. Extracellular vesicle fusion visualized by cryo-electron microscopy. PNAS Nexus 1, pgac156 (2022). 10.1093/pnasnexus/pgac156

24 Pinto da Silva, P. & Nogueira, M. L. Membrane fusion during secretion. A hypothesis based on electron microscope observation of Phytophthora Palmivora zoospores during encystment. J Cell Biol 73, 161–181 (1977). 10.1083/jcb.73.1.161

25 Zhao, W. D. et al. Hemi-fused structure mediates and controls fusion and fission in live cells. Nature 534, 548–552 (2016). 10.1038/nature18598

26 Luzio, J. P., Hackmann, Y., Dieckmann, N. M. & Griffiths, G. M. The biogenesis of lysosomes and lysosome-related organelles. Cold Spring Harb Perspect Biol 6, a016840 (2014). 10.1101/cshperspect.a016840

27 Bonifacino, J. S. & Glick, B. S. The mechanisms of vesicle budding and fusion. Cell 116, 153–166 (2004). 10.1016/s0092-8674(03)01079-1

28 Shvarev, D. et al. Structure of the HOPS tethering complex, a lysosomal membrane fusion machinery. Elife 11 (2022). 10.7554/eLife.80901

29 Luzio, J. P., Parkinson, M. D., Gray, S. R. & Bright, N. A. The delivery of endocytosed cargo to lysosomes. Biochem Soc Trans 37, 1019–1021 (2009). 10.1042/BST0371019

30 Brown, M. F. Curvature forces in membrane lipid-protein interactions. Biochemistry 51, 9782–9795 (2012). 10.1021/bi301332v

31 Callan-Jones, A., Sorre, B. & Bassereau, P. Curvature-driven lipid sorting in biomembranes. Cold Spring Harb Perspect Biol 3 (2011). 10.1101/cshperspect.a004648

32 Lipowsky, R. Remodeling of Membrane Shape and Topology by Curvature Elasticity and Membrane Tension. Adv Biol (Weinh) 6, e2101020 (2022). 10.1002/adbi.202101020

33 Chanturiya, A., Chernomordik, L. V. & Zimmerberg, J. Flickering fusion pores comparable with initial exocytotic pores occur in protein-free phospholipid bilayers. Proc Natl Acad Sci U S A 94, 14423–14428 (1997). 10.1073/pnas.94.26.14423

34 Risselada, H. J., Bubnis, G. & Grubmuller, H. Expansion of the fusion stalk and its implication for biological membrane fusion. Proc Natl Acad Sci U S A 111, 11043–11048 (2014). 10.1073/pnas.1323221111

35 Mattila, J. P. et al. A hemi-fission intermediate links two mechanistically distinct stages of membrane fission. Nature 524, 109–113 (2015). 10.1038/nature14509

36 Jahn, R., Lang, T. & Sudhof, T. C. Membrane fusion. Cell 112, 519–533 (2003). 10.1016/s0092-8674(03)00112-0

37 Cesar-Silva, D., Pereira-Dutra, F. S., Moraes Giannini, A. L. & Jacques, G. d. A. C. The Endolysosomal System: The Acid Test for SARS-CoV-2. Int J Mol Sci 23 (2022). 10.3390/ijms23094576

38 Gould, G. W. & Lippincott-Schwartz, J. New roles for endosomes: from vesicular carriers to multi-purpose platforms. Nat Rev Mol Cell Biol 10, 287–292 (2009). 10.1038/nrm2652

39 Navaroli, D. M. et al. Rabenosyn-5 defines the fate of the transferrin receptor following clathrin-mediated endocytosis. Proc Natl Acad Sci U S A 109, E471–480 (2012). 10.1073/pnas.1115495109

40 Hanson, P. I. & Cashikar, A. Multivesicular body morphogenesis. Annu Rev Cell Dev Biol 28, 337–362 (2012). 10.1146/annurev-cellbio-092910-154152

41 Jeger, J. L. Endosomes, lysosomes, and the role of endosomal and lysosomal biogenesis in cancer development. Mol Biol Rep 47, 9801–9810 (2020). 10.1007/s11033-020-05993-4

42 Ganesan, D. & Cai, Q. Understanding amphisomes. Biochem J 478, 1959–1976 (2021). 10.1042/BCJ20200917

43 Shimizu, H., Hosseini-Alghaderi, S., Woodcock, S. A. & Baron, M. Alternative mechanisms of Notch activation by partitioning into distinct endosomal domains. J Cell Biol 223 (2024). 10.1083/jcb.202211041

44 Lucic, V., Forster, F. & Baumeister, W. Structural studies by electron tomography: from cells to molecules. Annu Rev Biochem 74, 833–865 (2005). 10.1146/annurev.biochem.73.011303.074112

45 Arribas Perez, M. & Beales, P. A. Biomimetic Curvature and Tension-Driven Membrane Fusion Induced by Silica Nanoparticles. Langmuir 37, 13917–13931 (2021). 10.1021/acs.langmuir.1c02492

46 Woodall, N. B., Yin, Y. & Bowie, J. U. Dual-topology insertion of a dual-topology membrane protein. Nat Commun 6, 8099 (2015). 10.1038/ncomms9099

47 Vitrac, H., Dowhan, W. & Bogdanov, M. Effects of mixed proximal and distal topogenic signals on the topological sensitivity of a membrane protein to the lipid environment. Biochim Biophys Acta Biomembr 1859, 1291–1300 (2017). 10.1016/j.bbamem.2017.04.010

48 Herrmann, E., Langemeyer, L., Auffarth, K., Ungermann, C. & Kummel, D. Targeting of the Mon1-Ccz1 Rab guanine nucleotide exchange factor to distinct organelles by a synergistic protein and lipid code. J Biol Chem 299, 102915 (2023). 10.1016/j.jbc.2023.102915

49 Shirane, M. Lipid Transfer-Dependent Endosome Maturation Mediated by Protrudin and PDZD8 in Neurons. Front Cell Dev Biol 8, 615600 (2020). 10.3389/fcell.2020.615600

50 Bissig, C. & Gruenberg, J. Lipid sorting and multivesicular endosome biogenesis. Cold Spring Harb Perspect Biol 5, a016816 (2013). 10.1101/cshperspect.a016816

51 Mendoza, A. D. et al. Lysosome-related organelles contain an expansion compartment that mediates delivery of zinc transporters to promote homeostasis. Proc Natl Acad Sci U S A 121, e2307143121 (2024). 10.1073/pnas.2307143121

52 Smith, S. M. & Smith, C. J. Capturing the mechanics of clathrin-mediated endocytosis. Curr Opin Struct Biol 75, 102427 (2022). 10.1016/j.sbi.2022.102427

53 Sochacki, K. A. et al. The structure and spontaneous curvature of clathrin lattices at the plasma membrane. Dev Cell 56, 1131–1146 e1133 (2021). 10.1016/j.devcel.2021.03.017

54 Zhao, J. et al. Multiple claudin-claudin cis interfaces are required for tight junction strand formation and inherent flexibility. Commun Biol 1, 50 (2018). 10.1038/s42003-018-0051-5

55 Van Itallie, C. M., Tietgens, A. J. & Anderson, J. M. Visualizing the dynamic coupling of claudin strands to the actin cytoskeleton through ZO-1. Mol Biol Cell 28, 524–534 (2017). 10.1091/mbc.E16-10-0698

56 Iancu, C. V. et al. Electron cryotomography sample preparation using the Vitrobot. Nat Protoc 1, 2813–2819 (2006). 10.1038/nprot.2006.432

57 Hagen, W. J. H., Wan, W. & Briggs, J. A. G. Implementation of a cryo-electron tomography tilt-scheme optimized for high resolution subtomogram averaging. J Struct Biol 197, 191–198 (2017). 10.1016/j.jsb.2016.06.007

58 Kremer, J. R., Mastronarde, D. N. & McIntosh, J. R. Computer visualization of three-dimensional image data using IMOD. J Struct Biol 116, 71–76 (1996). 10.1006/jsbi.1996.0013

59 Zheng, S. et al. AreTomo: An integrated software package for automated marker-free, motion-corrected cryo-electron tomographic alignment and reconstruction. J Struct Biol X 6, 100068 (2022). 10.1016/j.yjsbx.2022.100068

60 Lamm, L. et al. MemBrain v2: an end-to-end tool for the analysis of membranes in cryo-electron tomography. bioRxiv, 2024.2001.2005.574336 (2024). 10.1101/2024.01.05.574336

61 Pettersen, E. F. et al. UCSF ChimeraX: Structure visualization for researchers, educators, and developers. Protein Science 30, 70–82 (2021). 10.1002/pro.3943

62 Stalling, D., Westerhoff, M. & Hege, H.-C. Amira: A highly interactive system for visual data analysis. The visualization handbook 38, 749–767 (2005).

